# SnapATAC2: a fast, scalable and versatile tool for analysis of single-cell omics data

**DOI:** 10.1101/2023.09.11.557221

**Authors:** Kai Zhang, Nathan R Zemke, Ethan J Armand, Bing Ren

## Abstract

Single-cell omics technologies have ushered in a new era for the study of dynamic gene regulation in complex tissues during development and disease pathogenesis. A major computational challenge in analyzing these datasets is to project the large-scale and high dimensional data into low-dimensional space while retaining the relative relationships between cells in order to decompose the cellular heterogeneity and reconstruct cell-type-specific gene regulatory programs. Conventional dimensionality reduction methods suffer from computational inefficiency, difficulty to capture the full spectrum of cellular heterogeneity, or inability to apply across diverse molecular modalities. Here, we report a fast and nonlinear dimensionality reduction algorithm that not only more accurately captures the heterogeneities of single-cell omics data, but also features runtime and memory usage that is computational efficient and linearly proportional to cell numbers. We implement this algorithm in a Python package named SnapATAC2, and demonstrate its superior performance, remarkable scalability and general adaptability using an array of single-cell omics data types, including single-cell ATAC-seq, single-cell RNA-seq, single-cell Hi-C, and single-cell multiomics datasets.

## Introduction

Rapid advancements in single-cell omics technologies have enabled the analysis of the gene regulatory programs encoded in the genome at unprecedented resolution and scale^1^. Single-cell analysis of genomes, transcriptomes, open chromatin landscapes, histone modifications, transcription factor binding, DNA methylation, chromatin architecture, *etc*., have provided valuable insights into the mechanisms governing cellular identity and regulation^1^. However, the extreme scale and complexity of single-cell omics data often present significant computational challenges, necessitating the development of efficient, scalable, and robust methods for data analysis^2^.

A crucial step in analyzing single-cell omics data is to project the high dimensional data onto low-dimensional space while retaining the relative relationships between cells, a process known as dimensionality reduction. This step is key to the success of downstream analyses such as clustering, batch correction, data integration, and visualization. Effective dimensionality reduction techniques are instrumental for visualization of distinct cell populations, identification of rare cell types, and delineation of cell-type-specific transcriptional regulatory programs^2^. Currently, single-cell omics dimensionality reduction algorithms fall into two main categories: linear and nonlinear techniques. Linear dimensionality reduction algorithms, such as principal component analysis (PCA), used by SCANPY^3^ and Seurat^4^, for single-cell RNA-seq (scRNA-seq) data analysis, and latent semantic indexing (LSI) used by ArchR^5^ and Signac^6^ for single-cell Assay for Transposase-Accessible Chromatin using sequencing (scATAC-seq) data analysis, are popular due to their computational efficiency and scalability.

However, these algorithms are not optimal for handling single-cell datasets with complex and nonlinear structures, such as single-cell Hi-C (scHi-C) and single-cell multimodal omics datasets. Nonlinear dimensionality reduction methods address these issues by more effectively capturing complex and often nonlinear cell relationships. Examples include latent Dirichlet allocation (LDA) used for scATAC-seq and scHi-C data^7,8^, Laplacian-based algorithms used for scRNA-seq and scATAC-seq data^9–13^, and various neural network models developed for scRNA-seq, scATAC-seq, and scHi-C data^14–18^. Nonlinear dimensionality reduction methods have also become the standard approach for single-cell data visualization. For example, t-distributed stochastic neighbor embedding (t-SNE)^19^ and uniform manifold approximation and projection (UMAP)^20^ are two widely-used algorithms for this purpose, despite recent concerns regarding their reliability and validity^21^. While nonlinear methods excel in handling complex structures and projecting data into low-dimensional manifolds, they are generally computationally inefficient, with limited scalability. For instance, LDA relies on the Markov chain Monte Carlo (MCMC) algorithm for model training, which is slow to converge, computationally expensive, and difficult to parallelize, making it difficult to be applied to large datasets^22^. Laplacian-based techniques like our previous work, SnapATAC^9^, necessitate computing similarity matrices between all pairs of cells, which leads to quadratic memory usage increase with the number of cells^23,24^. Deep neural network models, known for their high training costs, often require specialized computational hardware such as GPUs to be computationally feasible.

In the present study, we describe a nonlinear dimensionality reduction algorithm that achieves both computational efficiency and accuracy in discerning cellular composition of complex tissues from a broad spectrum of single-cell omics data types. The key innovation of our algorithm is the use of a matrix-free spectral embedding algorithm to project single-cell omics data into a low-dimensional space that preserves the intrinsic geometric properties of the underlying data. Unlike conventional spectral embedding approach that requires the construction of the graph Laplacian matrix, a process that demands a storage space increasing quadratically with the number of cells, our algorithm achieves the same goal while avoiding this computationally expensive step. Specifically, we utilize the Lanczos algorithm^25^ to derive eigenvectors while implicitly using the Laplacian matrix. This strategy significantly shortens the time and space complexity, making it linearly proportional to the number of the cells in the single-cell data. To evaluate the accuracy and utility of our algorithm, we conducted extensive benchmarking using a variety of datasets that encompass diverse experimental protocols, species, and tissue types. The results showed that our matrix-free spectral embedding algorithm outperforms existing methods in terms of speed, scalability and precision in resolving cell heterogeneity. Furthermore, we showed that our algorithm can be extended to diverse molecular modalities of single-cell omics datasets, revealing cell heterogeneity by leveraging complementary information from different single-cell omics data types. Lastly, we developed an approach capable of performing spectral embedding on data that exceeds available computer memory, a feature that will prove beneficial for atlas-scale single-cell omics datasets.

We have implemented these algorithmic advancements in a Python package called SnapATAC2. This package is a significant revamp of the original SnapATAC, offering substantial improvements such as increased speed, reduced memory usage, more reliable performance, and a comprehensive analysis framework for diverse single-cell omics data. SnapATAC2 is freely available at https://github.com/kaizhang/SnapATAC2.

## Results

### An overview of the SnapATAC2 workflow

SnapATAC2 is a comprehensive, high-performance solution for single-cell omics data analysis. Like the original SnapATAC^9^, SnapATAC2 streamlines end-to-end scATAC-seq data analysis with an extensive set of functionality. Moreover, SnapATAC2 is designed with flexibility in mind, intended for a variety of single-cell omics data types. For instance, its dimensionality reduction subroutine is readily applicable to scATAC-seq, scRNA-seq, single-cell DNA methylation, and scHi-C data, showcasing its adaptability. To enhance performance and scalability, SnapATAC2 uses the Rust^26^ programming language for executing computationally intensive subroutines, and provides a Python^27^ interface for seamless installation and user-friendly operation. This combination allows for efficient processing of large-scale single-cell omics data while maintaining accessibility for researchers across various levels of expertise. To further improve scalability when handling large-scale single-cell data, on-disk data structures and out-of-core algorithms are employed whenever possible. These modifications facilitate the analysis of large datasets without overburdening system resources. Additionally, SnapATAC2 is modular and adaptable, and allows users to tailor their analysis to specific requirements or integrate it with other software packages from the scverse^28^ ecosystem, such as SCANPY^3^ and scvi-tools^14^.

SnapATAC2 consists of four primary modules: preprocessing, embedding/clustering, functional enrichment analysis, and multi-modal omics analysis (Fig. 1a). Each module has been thoroughly revamped from the original SnapATAC, resulting in up to 100 times faster performance. The preprocessing module manages raw BAM files, calculates quality control metrics, generates cell-by-bin count matrices, and identifies doublets. These steps establish a robust foundation for subsequent analyses, ensuring data quality and integrity. At the heart of the SnapATAC2 package is the embedding/clustering module, featuring a novel matrix-free spectral embedding algorithm for dimensionality reduction, developed specifically in this study. This module incorporates dimensionality reduction, batch correction, and graph-based clustering, which are vital for distinguishing distinct cell populations and unveiling underlying biological structures. Functional enrichment analysis delves into various aspects of comprehensive data interpretation, including differential accessibility analysis and motif enrichment analysis. Lastly, the multi-modal analysis module enables the analysis of single-cell multimodal omics datasets, featuring more robust data integration, joint embedding of multiomic datasets, and the construction of gene regulatory networks utilizing information from multiple modalities.

**Figure 1.**
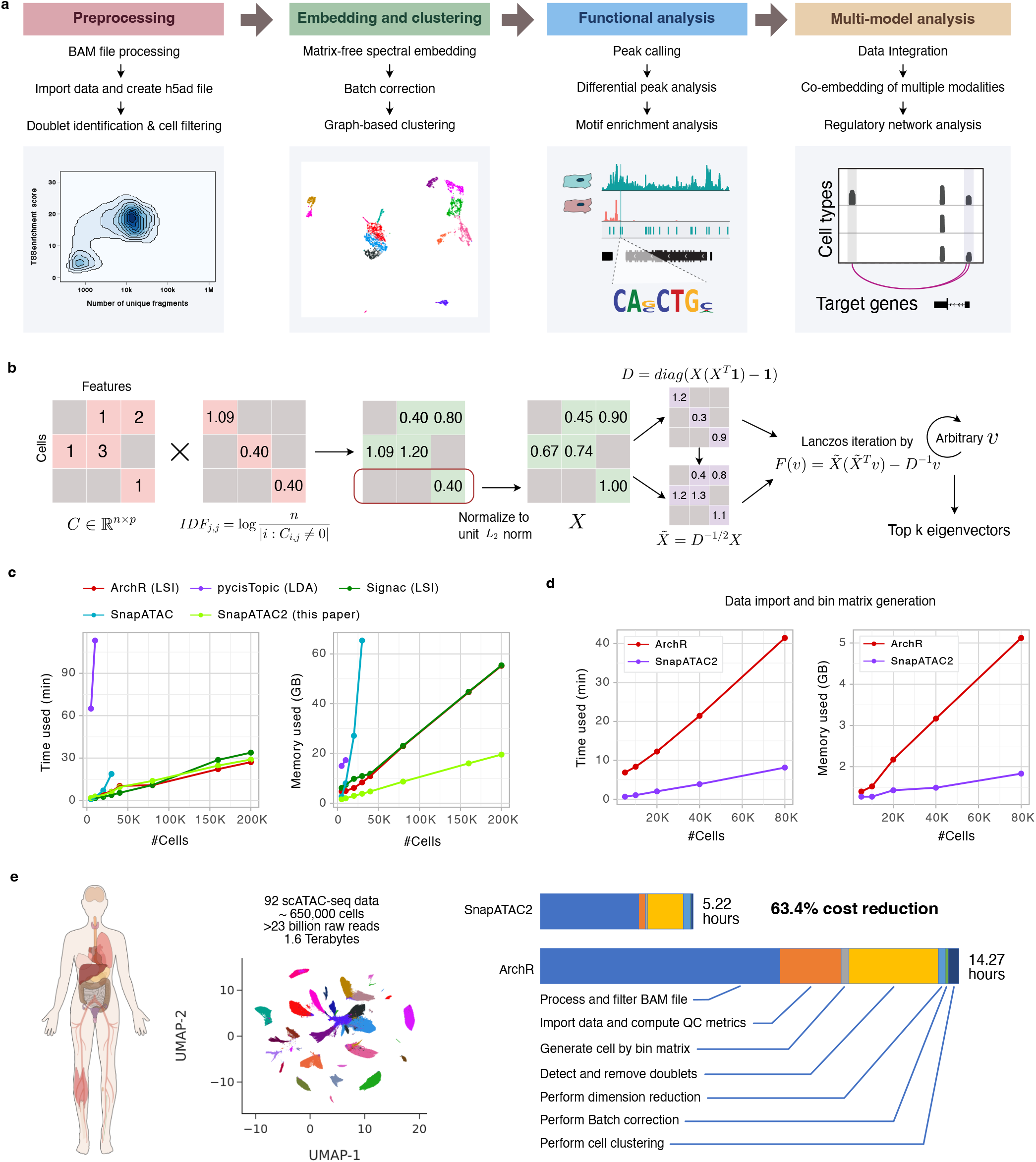
SnapATAC2 enables comprehensive and scalable end-to-end analysis of single-cell omics data. **a**, Overview of the SnapATAC2 Python package, featuring four primary modules: preprocessing, embedding/clustering, functional enrichment analysis, and multi-modal analysis. **b**, Schematic representation of the matrix-free spectral embedding algorithm in SnapATAC2, consisting of four main steps: feature scaling with inverse term frequency, row-wise *L*_2_ norm normalization, normalization using the degree matrix, and eigenvector calculation through the Lanczos algorithm^25^. **c**, Line plots comparing running times (left) and memory usage (right) of various dimensionality reduction algorithms for scATAC-seq data. Each data point indicates the average of three independent measurements. The benchmark was conducted on a Linux server with 4 cores and 120GB memory. See **Extended Data Table 1** for additional information. **d**, Line plots comparing running times (left) and memory usage (right) of SnapATAC2 and ArchR for preprocessing fragment files in scATAC-seq analysis. The benchmarked steps include fragment file processing, quality control metric computation, and cell-by-bin matrix generation. Each data point represents the average of three independent measurements. The benchmark was conducted on a standard Linux server with 4 cores and 120GB memory. See **Extended Data Table 2** for further details. **e**, Runtime comparison between ArchR and SnapATAC2 for end-to-end analysis of 92 raw bam files produced by scATAC-seq experiments. The benchmark was performed on a Linux server with 8 cores and 64GB memory.

### Matrix-free spectral embedding enables efficient and accurate dimensionality reduction of single-cell ATAC-seq data

Spectral embedding, also known as Laplacian eigenmaps, is a widely used technique for nonlinear dimensionality reduction^29^. This method boasts several key advantages, such as locality preservation, noise reduction, and a natural connection to clustering^29^. Spectral embedding techniques leverage the spectrum (eigenvalues and eigenvectors) of the cell similarity matrix calculated from single-cell omics datasets to perform dimensionality reduction. However, the computation of this matrix is a rate-limiting step and a memory bottleneck, creating challenges for handling datasets consisting of large numbers of cells. For example, the memory usage of the similarity matrix for a dataset with one million cells is approximately seven terabytes, far beyond the capacity of most computational servers. To address this barrier, we devise a matrix-free spectral embedding algorithm that efficiently computes eigenvectors using the Lanczos algorithm^25^, eliminating the need for constructing a full similarity matrix (Fig. 1b, see Methods for details). This method exhibits linear space and time usage relative to the input matrix size, resulting in a faster and memory-efficient approach for processing of large datasets. Notably, our algorithm avoids heuristic approximations, delivering precise solutions, distinguishing it from previous methods that generate approximate outcomes^10,11,30^ (see Methods for details).

To benchmark the performance of SnapATAC2, we generated synthetic scATAC-seq datasets with varying cell numbers and compared the scalability of the matrix-free spectral embedding algorithm to other widely-used dimensionality reduction algorithms, such as LSI (employed by ArchR^5^ and Signac^6^), LDA (used by cisTopic^7^), and classic spectral embedding with the Jaccard index (implemented in the original SnapATAC^9^ package). We excluded neural network-based methods from this comparison due to their significantly longer runtime and the requirement for specialized computer hardware. For example, training the scBasset^16^ neural network model on a dataset with 10,000 cells using 16 CPU cores takes roughly 10 minutes per epoch. With 500 epochs needed for training, the entire process would require 83 hours. In contrast, SnapATAC2 and ArchR can complete the analysis in under three minutes. As shown in Fig. 1c, the runtime of SnapATAC2’s matrix-free spectral embedding algorithm scales linearly with the number of cells, rivaling the fastest algorithm, LSI, which utilizes truncated singular value decomposition (SVD) for dimensionality reduction. In contrast, the original SnapATAC package experienced out-of-memory errors when processing datasets with over 40,000 cells on a Linux server equipped with 120 gigabytes of memory. Although cisTopic did not face memory constraints, it required approximately 2 hours to complete the analysis of 10,000 cells while SnapATAC2 finished in under 3 minutes. Notably, the SnapATAC2 package consumes the least amount of memory among all tested methods.

One of the objectives of SnapATAC2 is to offer comprehensive end-to-end analysis for scATAC-seq data. ArchR was previously considered the most scalable and complete software package for this purpose^31^. We therefore carried out a thorough comparison between SnapATAC2 and ArchR across eight stages in scATAC-seq analysis, including BAM file filtering and processing, data import, quality control metric computation, cell-by-bin matrix generation, doublet removal, dimensionality reduction, batch correction, and clustering. In Fig. 1d, we showed that SnapATAC2 significantly outpaces ArchR in preprocessing steps while maintaining sub-linear memory usage relative to the total number of cells analyzed. Subsequently, we conducted an end-to-end analysis using both SnapATAC2 and ArchR on our previously published human single-cell atlas of chromatin accessibility^32^. This atlas comprises 92 scATAC-seq samples, roughly 650,000 cells, over 23 billion raw reads, and has a total size of 1.6 terabytes. As shown in Fig. 1e, SnapATAC2 completed the entire process in 5.22 hours, whereas ArchR took 14.27 hours. At this scale, SnapATAC2 is almost three times faster than ArchR, decreasing computational costs by approximately 63.4%.

### SnapATAC2 produces the highest-quality cell embeddings among existing methods

We proceeded to assess the precision of our dimensionality reduction algorithm in identifying the relationships between cells, in comparison to other existing methods. For this purpose, we utilized a previously published benchmark dataset of synthetic scATAC-seq data^22^, consisting of ten simulated datasets with varying sequencing depths and noise levels, along with ground-truth cell type labels (Fig. 2a). To increase the difficulty of this task, we combined all ten datasets into a single dataset. We expanded our comparison to include deep neural network-based algorithms including PeakVI^15^ and scBasset^16^, as they have been reported to surpass traditional dimensionality reduction methods. We also included EpiScanpy^33^, a method that utilizes PCA for dimensionality reduction. In total, we benchmarked eight methods on the merged dataset, as shown in Fig. 2b. After dimensionality reduction, we applied graph-based clustering using the Leiden algorithm^34^ and assessed the clustering quality with the adjusted Rand index (ARI) and adjusted mutual information (AMI). ARI and AMI are both measures to compare the similarity between two data clusterings and have been routinely employed to assess the performance of clustering algorithms^22,35^. We hypothesized that high-quality embeddings should yield clusters consistent with the ground-truth cell type labels, and hence resulted in high ARI and AMI scores. In Fig. 2b,c, we showed that SnapATAC2 and the original SnapATAC produced the highest ARI and AMI scores, followed by scBasset and LDA (cisTopic). On the contrary, ArchR and PeakVI generated the poorest embeddings, with significantly lower ARI and AMI scores than other methods, likely because these methods were not designed for handling substantial noise levels that were purposely introduced in this benchmark. These results show that the original SnapATAC and our newly developed SnapATAC2 are the only methods that are robust to large noise levels and produce high-quality embeddings.

**Figure 2.**
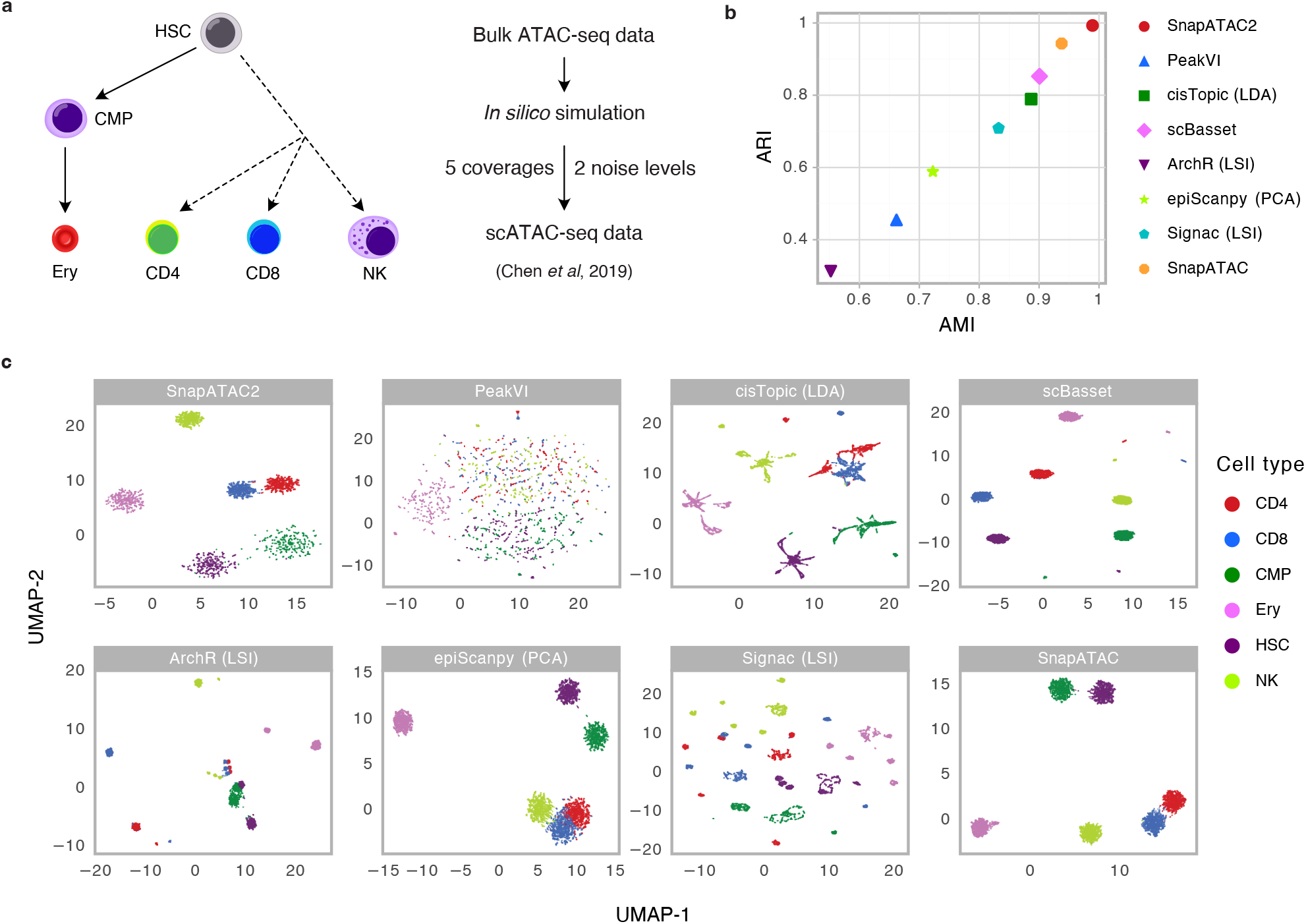
Benchmarking of SnapATAT2 and other dimensionality reduction algorithms using synthetic scATAC-seq data. **a**, Synthetic scATAC-seq datasets^22^ with different sequencing depths (5000, 2500, 1000, 500, and 250 fragments per cell) and noise levels (0, 0.2, and 0.4) were merged into a single dataset. The merged dataset contains 9,600 cells and includes the following six cell types: hematopoietic stem cells (HSCs), common myeloid progenitor cells (CMPs), erythroid cells (Ery), natural killer cells (NK), CD4^+^ T cells (CD4), and CD8^+^ T cells (CD8). **b**, Dot plot comparing the Adjusted Rand Index (ARI, Y-axis) and adjusted mutual information (AMI, X-axis) for eight dimensionality reduction methods. Refer to **Extended Data Table 3** and **Fig. S1** for further details and additional method comparisons. **c**, UMAP visualization of the embeddings generated by each method for the simulated dataset. Individual cells are color-coded based on the cell type labels indicated in **a**.

To assess SnapATAC2’s performance under conditions that mimic data generated from real experiments, we utilized a previously published scATAC-seq dataset from the human hematopoietic system^36^. This dataset comprises 2,034 hematopoietic cells that were profiled and sorted using fluorescence-activated cell sorting (FACS) from ten cell populations, including hematopoietic stem cells (HSCs), multipotent progenitors (MPPs), lymphoid-primed multipotent progenitors (LMPPs), common myeloid progenitors (CMPs), granulocyte-macrophage progenitors (GMPs), GMP-like cells, megakaryocyte-erythroid progenitors (MEPs), common lymphoid progenitors (CLPs), monocytes (mono), and plasmacytoid dendritic cells (pDCs) (Fig. 3a). SnapATAC2 again achieved the highest ARI and AMI scores, followed by other nonlinear methods including cisTopic, PeakVI and scBasset (Fig. 3b,c). Overall, these two benchmarking experiments demonstrate that nonlinear dimensionality reduction algorithms generally outperform linear methods, and SnapATAC2 produces the highest-quality cell embeddings among all methods tested.

**Figure 3.**
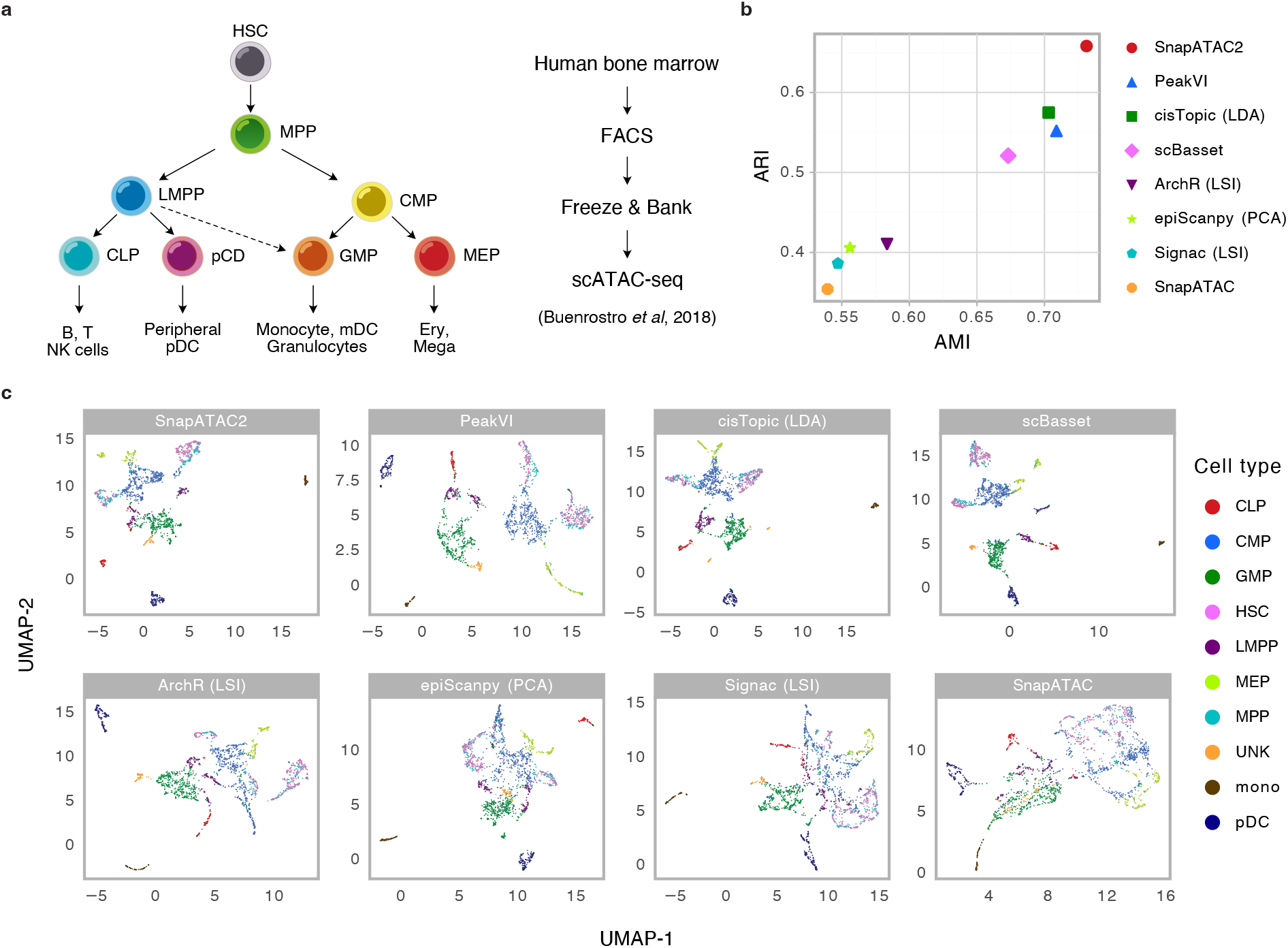
Benchmarking of SnapATAT2 and other dimensionality reduction algorithms using actual scATAC-seq data with ground-truth labels. **a**, Overview of cell types analyzed in the Buenrostro2018 scATAC-seq dataset, which includes 2,034 hematopoietic cells profiled and FACS-sorted from ten cell populations: hematopoietic stem cells (HSCs), multipotent progenitors (MPPs), lymphoid-primed multipotent progenitors (LMPPs), common myeloid progenitors (CMPs), granulocyte-macrophage progenitors (GMPs), GMP-like cells, megakaryocyte-erythroid progenitors (MEPs), common lymphoid progenitors (CLPs), monocytes (mono), and plasmacytoid dendritic cells (pDCs). **b**, Dot plot comparing the Adjusted Rand Index (ARI, Y-axis) and adjusted mutual information (AMI, X-axis) for eight dimensionality reduction methods. Refer to **Extended Data Table 3** and **Fig. S2** for further details and additional method comparisons. **c**, UMAP visualization of the embeddings generated by each method for the Buenrostro2018 dataset. Individual cells are color-coded based on the cell type labels indicated in **a**.

Although SnapATAC2 achieved top performance in both benchmarking experiments, the two benchmark datasets have limited scope and may not fully represent the diversity of scATAC-seq data. Consequently, we examined additional public scATAC-seq datasets generated using different technologies, species, and tissue types^37–42^ (Table 1). Acknowledging the potential variations in cell type label quality between datasets, we restricted our analysis to broadly cited datasets. Additionally, we included six datasets where cells were annotated by an independent modality, rather than the ATAC modality, to minimize potential biases from the algorithm used to generate the cell type labels. SnapATAC2 achieved the highest AMI scores in six of the nine tested datasets and ranked among the top three methods in the remaining datasets (Fig. 4a). ARI scores exhibited similar trends, with SnapATAC2 achieving the highest scores in five of the nine datasets (Fig. 4b). In general, nonlinear dimensionality reduction methods, including SnapATAC2, PeakVI, scBasset, and cisTopic, outperformed linear methods such as EpiScanpy, ArchR, and Signac in resolving cellular heterogeneity in scATAC-seq data. The median rank of SnapATAC2 across all eleven datasets was 1, followed by PeakVI and scBasset, both with a median rank of 3 (Fig. 4c,d). Beyond its superior accuracy in identifying cell types, SnapATAC2 offers key advantages over deep neural network-based methods. It can operate without specialized hardware, consume significantly less runtime, maintain consistent performance across various datasets, and does not require extensive hyperparameter tuning.

**Table 1.**
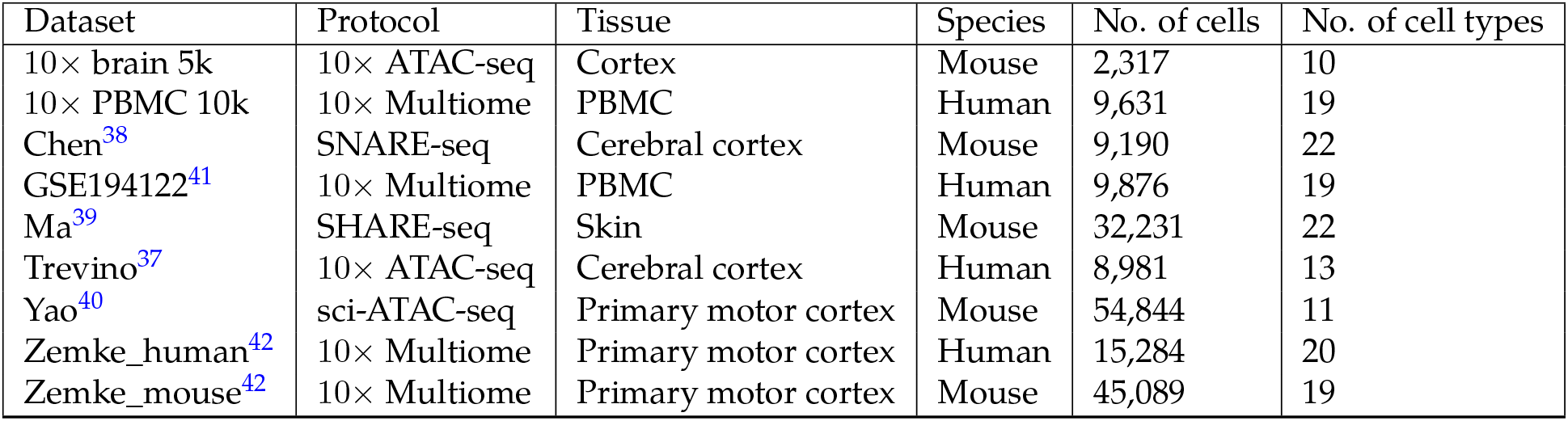
Curated scATAC-seq benchmark datasets used in the present study.

| Dataset | Protocol | Tissue | Species | No. of cells | No. of cell types |
| --- | --- | --- | --- | --- | --- |
| 10× brain 5k | 10× ATAC-seq | Cortex | Mouse | 2,317 | 10 |
| 10× PBMC 10k | 10× Multiome | PBMC | Human | 9,631 | 19 |
| Chen<sup>38</sup> | SNARE-seq | Cerebral cortex | Mouse | 9,190 | 22 |
| GSE194122<sup>41</sup> | 10× Multiome | PBMC | Human | 9,876 | 19 |
| Ma<sup>39</sup> | SHARE-seq | Skin | Mouse | 32,231 | 22 |
| Trevino<sup>37</sup> | 10× ATAC-seq | Cerebral cortex | Human | 8,981 | 13 |
| Yao<sup>40</sup> | sci-ATAC-seq | Primary motor cortex | Mouse | 54,844 | 11 |
| Zemke_human<sup>42</sup> | 10× Multiome | Primary motor cortex | Human | 15,284 | 20 |
| Zemke_mouse<sup>42</sup> | 10× Multiome | Primary motor cortex | Mouse | 45,089 | 19 |

**Figure 4.**
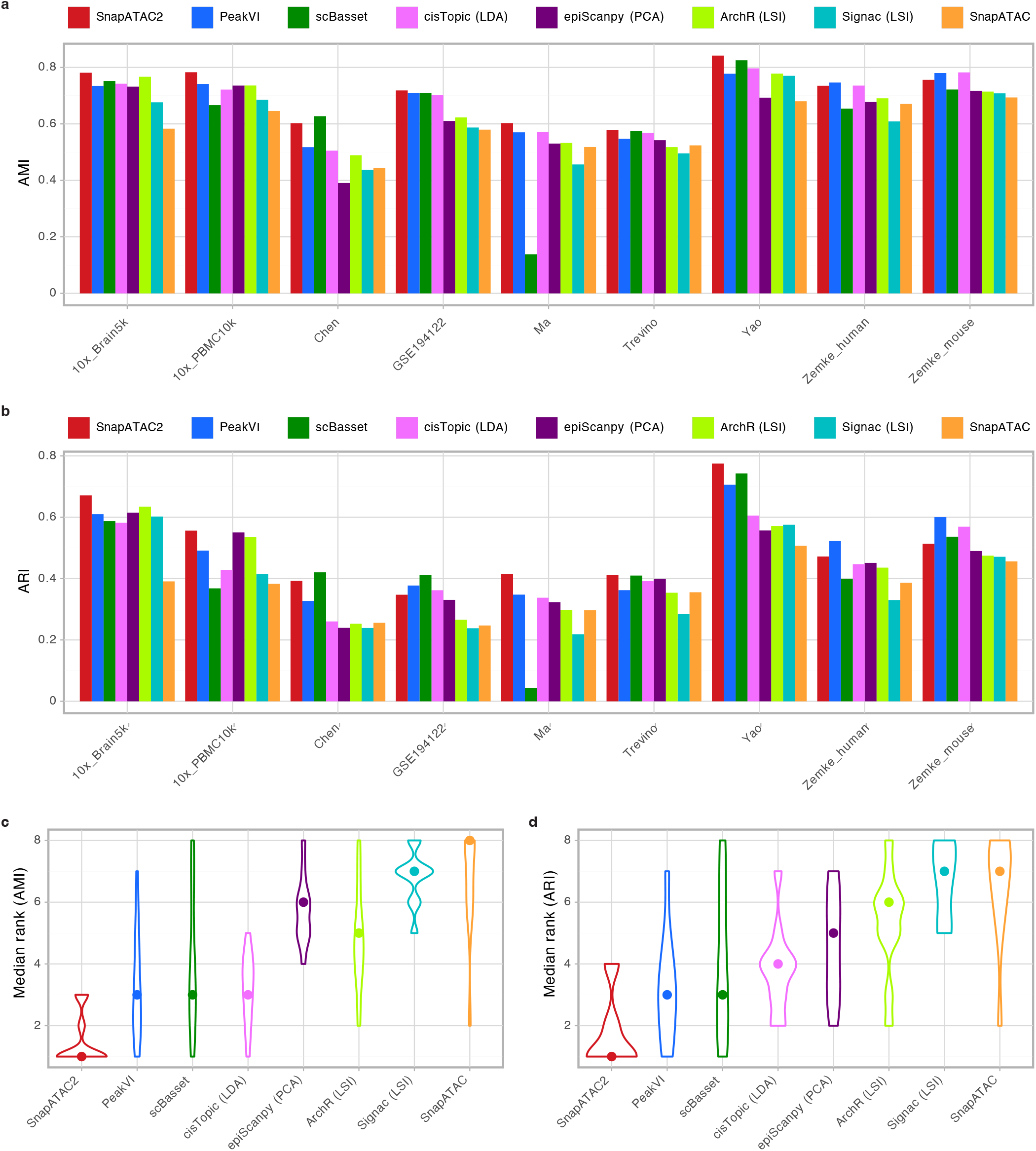
SnapATAC2 demonstrates superior performance over other methods on a wide range of datasets. **a**,**b**, Bar plots illustrating the AMI (**a**) and ARI (**b**) scores of eight methods on nine previously annotated scATAC-seq datasets. For more information, refer to **Extended Data Table 3** and **Figs. S3-S12. c**,**d**, Violin plots showing the distribution of AMI (**c**) and ARI (**d**) scores for each method across all eleven scATAC-seq benchmark datasets. The median rank of each method is indicated by the dot in the violin plot.

### Application of matrix-free spectral embedding to multiple single-cell omics data types

Spectral embedding has proven to be a versatile and effective technique across a broad spectrum of applications. We next explored whether this algorithm could be applied to other single-cell data types, such as scRNA-seq and scHi-C.

scHi-C data is notably sparse and exhibits an extraordinarily high dimensionality. Current computational methods struggle to fully utilize sparse scHi-C data for analyzing cell-to-cell variability in 3D genome features. Therefore, we initially focused on scHi-C data and tested our method, SnapATAC2, on two datasets with multiple cell types or known cell state information, including a sci-Hi-C dataset^8^ made public by the 4D Nucleome Project (4DN) and a dataset from Lee *et al*.^43^. We converted scHi-C data into a cell-by-feature count matrix by flattening the contact matrices of individual cells into vectors. This count matrix served as input for SnapATAC2’s matrix-free spectral embedding algorithm. The resulting cell embeddings exhibited clear patterns corresponding to the underlying cell types and cellular states (Fig. 5a,b, Fig. S13). We compared the quality of cell embeddings generated by SnapATAC2 with two existing methods: Higashi^18^ and scHiCluster^44^. SnapATAC2 significantly outperformed scHiCluster on both datasets and demonstrated comparable performance to Higashi, the current state-of-the-art method for scHi-C analysis. In the 4DN sci-Hi-C dataset, SnapATAC2 slightly outperformed Higashi, with an ARI score of 0.925 compared to 0.874 (Fig. 5a,b). In the Lee et al. dataset, Higashi (ARI: 0.721) slightly surpassed SnapATAC2 (ARI: 0.639) in identifying specific cell populations (Fig. S13). However, compared to Higashi, SnapATAC2 offers a significantly reduced runtime (Extended Data Table 4) and does not require specialized hardware, making it a more practical solution for analyzing large-scale scHi-C datasets.

**Figure 5.**
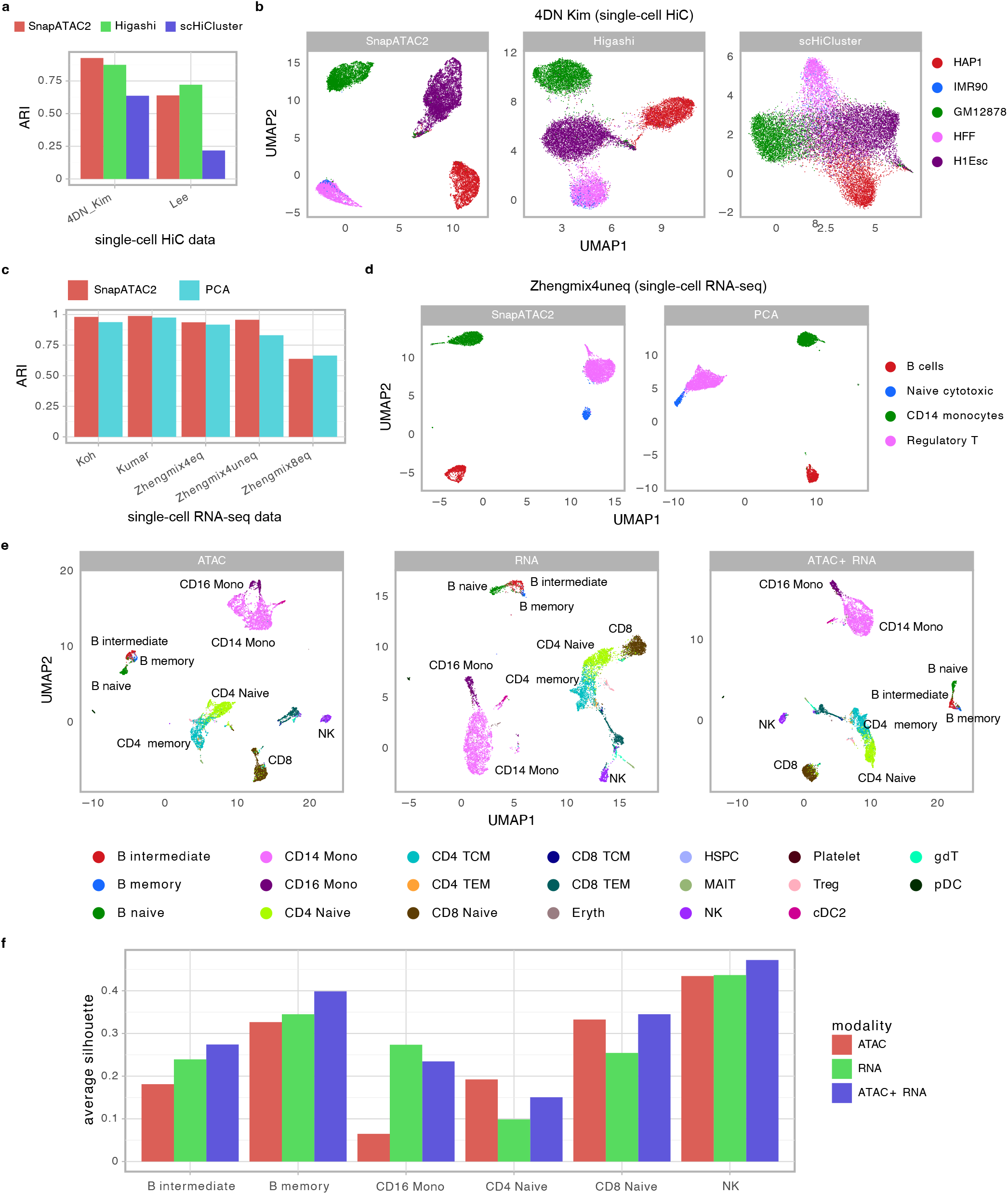
SnapATAC2 enables analysis of diverse single-cell omics data types. **a**, Bar plot comparing SnapATAC2, Higashi, and scHiCluster in terms of clustering performance for scHi-C data. For more details, see **Extended Data Table 4. b**, UMAP visualization of the embeddings generated by SnapATAC2, Higashi, and scHiCluster for the 4DN Kim dataset^8^. Cells are color-coded based on cell type labels. **c**, Bar plot comparing SnapATAC2 and PCA (implemented in SCANPY) for clustering scRNA-seq datasets^35^. For more details, see **Extended Data Table 5. d**, UMAP visualization of the embeddings produced by SnapATAC2 and PCA (SCANPY) for the Zhengmix4uneq dataset^35^. Cells are color-coded according to cell type labels. **e**, UMAP visualization of the embeddings generated by SnapATAC2 using ATAC modality (left panel), RNA modality (middle panel), or both modalities (right panel). Cells are color-coded based on cell type labels. **f**, Bar plot comparing the average silhouette scores of selected cell types derived from embeddings produced by ATAC modality, RNA modality, or both modalities.

We also applied SnapATAC2 to scRNA-seq datasets. PCA is commonly used in scRNA-seq analysis pipelines, such as Seurat^4^ and SCANPY^3^, and has been shown to be highly effective in identifying cell types^35,45^. In four out of five scRNA-seq benchmark datasets^35^, SnapATAC2 outperformed PCA, yielding embeddings more consistent with the underlying cell types (Fig. 5c,d and Fig. S14). Overall, SnapATAC2 slightly surpassed PCA, achieving an average ARI score of 0.900, while PCA scored 0.866. Notably, SnapATAC2 does not require data centering, a common preprocessing step in PCA analysis. When running PCA on uncentered data, the average ARI dropped to 0.841 (Extended Data Table 5), indicating that the centering step is crucial for optimal PCA performance. However, centering effectively transforms a sparse matrix into a dense matrix, which can be computationally expensive and prohibitive for large-scale datasets.

SnapATAC2’s dimensionality reduction algorithm is also applicable to single-cell DNA methylation data. When applied to a 5-methylcytosine sequencing 2 (snmC-seq2) data generated in mouse pituitaries^46^, SnapATAC2 produced cell embeddings that are largely consistent with the cell types identified by the original study (Fig. S15). Notably, the method provided finer resolution for some cell types, such as somatotropes and lactotropes.

In conclusion, SnapATAC2 is a versatile and effective method for the analysis of various single-cell data types, including scATAC-seq, scHi-C, scRNA-seq, and single-cell DNA methylation data. It demonstrates comparable or superior performance to existing methods, while offering practical advantages such as reduced runtime and no need for specialized hardware.

### SnapATAC2 enables robust joint embedding of single-cell multiomics data

The rapid expansion of single-cell multimodal omics technologies, such as 10× multiome (ATAC/RNA-seq), Paired-Tag^47^, and single-cell Methyl-HiC/snm3C-seq^43^, has provided powerful tools for investigating gene regulatory mechanisms. However, current methods for analyzing these data often focus on a single modality, underutilizing the full potential of multimodal datasets. We therefore investigated the applicability of our algorithm to single-cell multimodal omics data. Multi-view spectral embedding is an extension of spectral embedding, which enables the joint embedding of multiple data representation views. This method has demonstrated its ability to harness complementary information from individual views and enhance performance in downstream analyses, making it an ideal candidate for analyzing single-cell multiomics data. The multi-view spectral embedding process typically consists of three steps: first, a similarity or kernel matrix is calculated from each view; second, a joint kernel matrix is constructed by combining or co-regularizing the kernel matrices in a certain manner; and lastly, spectral embedding is performed using the joint kernel matrix. In this study, we opted for kernel addition to combine the kernel matrices, as it has proven to be an effective method for achieving excellent clustering results^47,48^. Moreover, kernel addition enables the extension of the matrix-free spectral embedding algorithm to multi-view spectral embedding while maintaining the linear time and space complexity of the algorithm (see Methods for details).

We applied this matrix-free multi-view spectral embedding algorithm to a 10× Genomics multiome dataset, which jointly profiles chromatin accessibility and the transcriptome for approximately 10,000 human peripheral blood mononuclear cells (PBMCs). To better evaluate the algorithm’s performance, we first annotated the cells according to a previously published single-cell atlas of human PBMCs^4^. To compare the performance of joint embedding with individual views, we also performed spectral embedding on each modality separately. Our findings revealed that, while independent unsupervised analyses of RNA and ATAC data generated predominantly consistent cell classifications, there were notable differences present (Fig. 5e,f). For instance, CD8^+^ and CD4^+^ T cells were close to each other when analyzing the transcriptome, but separated clearly in the ATAC data (Fig. 5e,f). Conversely, intermediate and memory B cells partially overlapped when analyzing the ATAC data but were more distinguishable in the transcriptomic data (Fig. 5e,f). In comparison to the separate analysis of either modality, multi-view spectral embedding using both modalities clearly separated CD4^+^ and CD8^+^ T cells and uncovered subtle heterogeneity within B cells. Overall, the joint embedding of ATAC and RNA data enhanced the separation of cell types and revealed subtle heterogeneity within cell types, as evidenced by the increased silhouette scores across different cell types (Fig. 5f).

### SnapATAC2 overcomes memory constraints for analyzing atlas-scale datasets

SnapATAC2’s matrix-free spectral embedding algorithm features linear space and time complexity, positioning it among the top-performing solutions. With 120 gigabytes of memory, SnapATAC2 can effectively perform dimensionality reduction for millions of cells. However, memory may still be a limiting factor when processing atlas-scale datasets, especially when combining data from multiple consortium studies or analyzing those with extremely high sequencing depth. To address this limitation, we developed an algorithm that circumvents the memory bottleneck of a computer, enabling the processing of massive numbers of cells in a reasonable timeframe (Fig. 6a). Our approach involved creating an on-disk data structure for efficient storage and access to portions of the count matrix without loading the entire matrix into memory. We then performed matrix-free spectral embedding on a subsampled count matrix to obtain a reference embedding. Using this reference embedding, we applied the Nyström algorithm^49^ to calculate the embedding for all cells without needing to ever store the complete count matrix in memory. Additionally, the final process can be distributed across multiple machines for a high degree of parallelism, enabling the processing of billions of cells relatively quickly. This feature is essential for analyzing large amounts of single-cell data generated from consortium efforts such as the BRAIN Initiative Cell Atlas Network (BICAN), the Human Cell Atlas (HCA), and the Human BioMolecular Atlas Program (HuBMAP).

**Figure 6.**
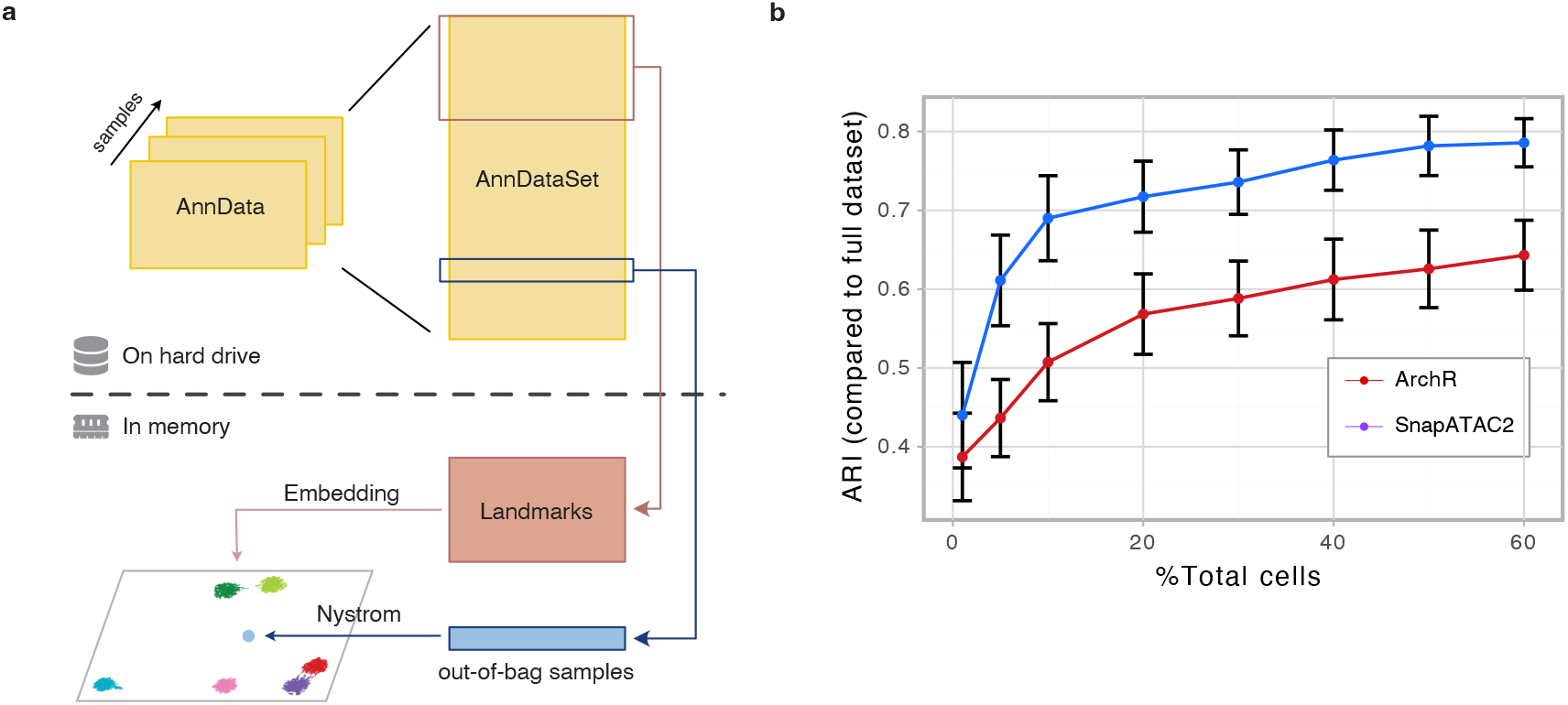
SnapATAC2 overcomes memory constraints for analyzing atlas-scale datasets. **a**, Schematic representation of the out-of-core algorithm implemented in SnapATAC2 for processing datasets that surpass available memory capacity. **b**, Line plot comparing ARI scores when undergoing different levels of subsampling. ARI scores were calculated by comparing the embeddings generated from the complete dataset with those predicted using the subsampled dataset. Error bars represent standard deviation of measurements across nine independent datasets. See **Extended Data Table 6** for further details.

Subsampling and approximation inherently lead to a reduction in accuracy. To evaluate this effect, we compared the embeddings generated from a subsampled dataset to those produced by the complete dataset. We discovered that our out-of-core algorithm yielded embeddings that were largely consistent with those generated by the complete dataset at various subsampling rates (Fig. 6b and Extended Data Table 6). For comparison, we conducted the same analysis using the subsampled LSI algorithm implemented in ArchR. Our algorithm demonstrated greater accuracy across different subsampling rates compared to subsampled LSI (Fig. 6b). For instance, at a 10% sampling rate, the average ARI score across nine experiments for the embeddings generated by our algorithm was nearly 0.7, while the average ARI score for those generated by subsampled LSI was about 0.5 (Fig. 6b). This finding suggests that SnapATAC2 is more robust to subsampling approaches necessary for analyzing large-scale datasets.

## Discussion

In the present study, we describe a software package named SnapATAC2 for the analysis of a diverse array of single-cell omics data. The performance of SnapATAC2 exceeds that of existing dimensionality reduction methods in terms of accuracy, noise robustness, and scalability, thus providing researchers with a powerful tool for investigating gene regulatory programs using single-cell genomics, transcriptomics and epigenomics analysis.

The key innovation of SnapATAC2 lies in its matrix-free spectral embedding algorithm for dimensionality reduction. While numerous algorithms have been proposed to expedite spectral embedding^10,11,23,30^, our algorithm stands out as it does not rely on sub-sampling or approximations, delivering the exact solution. This algorithm not only outperforms current methods in identifying cell clusters and heterogeneity but also maintains computational efficiency, making it highly suitable for large-scale single-cell omics data analysis. Furthermore, we demonstrated the versatility of the matrix-free spectral embedding algorithm by applying it to various single-cell data types, including scATAC-seq, scRNA-seq, single-cell DNA methylation and scHi-C data. Additionally, the multi-view extension of this algorithm enables the joint embedding of single-cell multiomics data and better reveals cellular heterogeneity of a mixed population, which is essential for understanding the complex dynamics of cellular states and functions. This cross-modality capability suggests that the underlying algorithm of SnapATAC2 can be adapted for analyzing different single-cell data types, potentially serving as the foundation for a unified multi-omics framework.

One limitation of the matrix-free spectral embedding algorithm is that it currently is implemented using only cosine function based similarity. For some data types, researchers may prefer to use other metrics to quantify the cell-to-cell similarity. For instance, in our findings, the Euclidean distance yielded more accurate results for the protein expression data used in Cellular Indexing of Transcriptomes and Epitopes by Sequencing (CITE-seq) experiments^50^.

Future developments could extend the matrix-free algorithm to accommodate other similarity metrics. For instance, a potential solution involves leveraging a small set of landmark points to transform the given data into sparse feature vectors^51^, followed by the application of the scalable matrix-free spectral embedding algorithm. In conclusion, SnapATAC2 represents a substantial advancement in single-cell data analysis, offering an accessible, scalable, and high-performance solution for researchers studying epigenomics. With continued development and optimization, SnapATAC2 has the potential to become a general tool in single-cell multi-omics data analysis, ultimately facilitating novel biological discoveries.

## Methods

### SnapATAC2’s high performance preprocessing module

SnapATAC2’s preprocessing module including BAM file processing, fragment file importing, QC metrics calculation, and cell by feature count matrix generation. All functions in this module are implemented using out-of-core algorithms, which enables SnapATAC2 to handle large-scale single-cell datasets efficiently without being limited by memory constraints.

### Converting BAM files to fragment files

SnapATAC2 is capable of processing unfiltered BAM files, converting them into files containing the genomic coordinates of the fragments. A fragment record consists of five specific fields: the reference genome chromosome of the fragment, the adjusted start position of the fragment on the chromosome, the adjusted end position of the fragment on the chromosome, the cell barcode associated with the fragment, and the total number of read pairs linked to the fragment. In order to convert BAM files into fragment files, the following steps are taken. First, reads that are unmapped, not primary alignment, improperly aligned, have mapping quality less than 30, fail platform/vendor quality checks, or are optical duplicates are removed. Next, using out-of-core algorithms, the remaining reads are sorted by cell barcodes, and duplicate reads for each unique cell barcode are eliminated. Finally, the filtered and deduplicated BAM records are converted into fragments. This process is implemented by the “snapatac2.pp.make_fragment_file” function within the SnapATAC2 package.

### Compute QC metrics

SnapATAC2 offers a comprehensive set of QC metrics to assess data quality, which include the number of unique fragments per cell, the proportion of mitochondrial reads per cell, the proportion of duplicate reads per cell, and the TSS enrichment score per cell. To calculate the TSS enrichment score, we first retrieved the TSS positions from a gene annotation file. We then aggregated Tn5-corrected insertions within a +/-2000 bp range relative to each unique TSS (strand-corrected) across the entire genome. Subsequently, this profile was normalized based on the mean accessibility within the +/-(1900 to 2000) bp range from the TSS and smoothed at 11 bp intervals. The maximum value of the smoothed profile was taken as the TSS enrichment. The SnapATAC2 package’s “snapatac2.pp.import_data” function facilitates this process.

### Compute cell by feature count matrix

SnapATAC2 provides various methods to calculate the cell-by-feature count matrix. The “snapatac2.pp.add_tile_matrix” function computes the cell-by-bin matrix by dividing the genome into a fixed number of bins. The “snapatac2.pp.make_peak_matrix” function calculates the cell-by-peak matrix based on the peak file, while the “snapatac2.pp.make_gene_matrix” function computes the cell-by-gene matrix using the gene annotation file.

### Dimensionality reduction using spectral embedding

In this section we outline the core algorithms used to perform dimensionality reduction in the SnapATAC2 package. We first describe the preprocessing steps and then the classic spectral embedding method that works for arbitrary similarity metrics. Finally, we describe the matrix-free spectral embedding algorithm that works only for cosine similarity, but substantially decreases the running time and memory usage. Note the steps described below can be accomplished using the “snapatac2.tl.spectral” function from the SnapATAC2 package.

### Preprocessing

Given a cell by feature count matrix *C* ∈ ℝ^*n*×*p*^, we first scale the columns of the matrix by the Inverse Document Frequency (IDF). The IDF of a column or a feature *f* is defined by *idf* 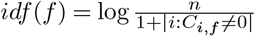

### Spectral embedding

Assuming the cell by feature count matrix *C* has been preprocessed according to the procedures described above, in classic spectral embedding we first compute the *n* × *n* pairwise similarity matrix *W* such that *W*_*ij*_ = *δ*(*C*_*i*∗_, *C*_*j*∗_), where *δ* : ℝ^*p*^ ×ℝ^*p*^ → ℝ is the function defines the similarity between any two cells. Typical choices of *δ* include the Jaccard index and the cosine similarity. We then compute the symmetric normalized graph Laplacian *L*_*sym*_ = *I* − *D*^−1*/*2^*WD*^−1*/*2^, where *I* is the identity matrix and *D* = *diag*(*W* 1). The bottom eigenvectors of *L*_*sym*_ are selected as the lower dimensional embedding. The corresponding eigenvectors can be computed alternatively as the top eigenvectors of the similarly normalized weight matrix: 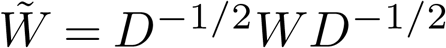

### Matrix-free spectral embedding with cosine similarity

In this section, we introduce a matrix-free algorithm for spectral embedding that avoids calculating the similarity matrix. This approach is specifically designed for cosine similarity. The cosine similarity between two vectors *A* and *B* is given by 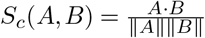 To express the cosine similarity using matrix operations, we first rescale the non-negative count matrix *C* to obtain a new matrix *X*, such that the rows of *X* have unit *L*_2_ norm. Consequently, the cosine similarity matrix between rows of *X* can be represented as *XX*^*T*^ .

In traditional spectral clustering algorithms, it is necessary to set the diagonals of the similarity matrix to zero^52^. This can be accomplished by subtracting the identity matrix from the similarity matrix, resulting in the final similarity matrix *W* = *XX*^*T*^ − *I*. The degree matrix can then be calculated as *D* = *diag*((*XX*^*T*^ − *I*)**1**) = *diag*(*X*(*X*^*T*^ **1**) − **1**).

The normalized similarity matrix, denoted as 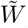 can then be computed as follows:

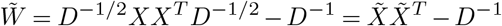

where 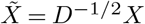 It is important to note that 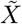 has the same dimensions as *X*, and if *X* is sparse, 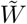preserves the sparsity pattern of *X*. Conventional spectral embedding algorithms compute 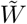 and select its top eigenvectors as the lower-dimensional embedding. Previous work has attempted to compute the top eigenvectors of an approximation of 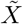to avoid the need for computing the full similarity matrix^30^. In other studies^10,11^, the authors chose not to set the diagonals of the similarity matrix to zero. Consequently, the eigendecomposition of 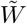 is equivalent to the SVD of 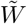 which can be computed efficiently. However, our benchmarking reveals that setting the diagonal of *W* to zero is necessary as it significantly improve the embedding quality (Fig. S12).

Unlike previous work, we offer an exact solution to the problem. We apply the Lanczos algorithm^25^, an iterative method for computing the top eigenvectors of a symmetric matrix, to our problem without ever calculating 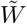 This requires computing the matrix-vector product between 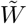 and **v** in each iteration, as follows: 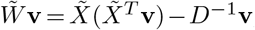where **v** is current solution to the eigenvalue problem and is iteratively refined by the Lanczos algorithm. By using the specific order of operations shown in the formula, we can reduce the computational cost of the matrix-vector product to 2*z* + *n*, where *n* is the number of rows in *X* and *z* is the number of non-zero elements in *X*. In comparison, performing this operation on the full similarity matrix requires *n*^2^ computations, which is prohibitively expensive for large number of cells. Thus, our matrix-free method is significantly faster and more memory efficient. The pseudocode for our algorithm is shown in Fig. S16a.

### Nyström method for out-of-sample embedding

The matrix-free method described above is very fast and memory efficient. However, for massive datasets with hundreds of millions of cells, storing the cell by feature count matrix *C* may already be a challenge. In this section, we describe an on-line embedding method that can be applied to datasets that do not fit into physical memory. The key idea here is to use the Nyström method^24^ to construct low-rank matrix approximations.

Given a symmetric similarity matrix *W* ∈ ℝ^*n*×*n*^, the Nyström method constructs a low-rank approximation 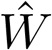 of *W* by sampling *l* ≪ *n* columns from *W*. Without loss of generality, we assume that the first *l* columns of *W* are sampled. The *W* matrix can then be partitioned into four blocks: 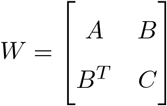, where columns 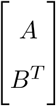 are our samples. The low-rank approximation 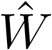 is then defined as,

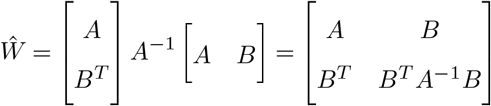

As a result, the degree matrix *D* can be approximated as,

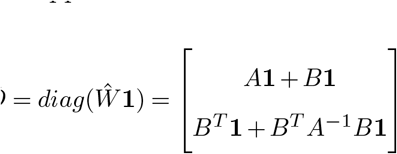

The eigenvector of 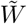 can then be approximated by,

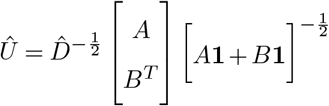

The time complexity of the approach discussed above is 𝒪(*nl*^2^ + *l*^3^). The space complexity amounts to 𝒪(*nl*), as it is necessary to store the sampled columns of *W*. In large-scale applications, sampling even a small subset of columns can prove to be exceedingly costly. As a result, rather than computing the inverse of *A*, we utilize an algorithm previously described^49^ to approximate the degree matrix. The full algorithm’s pseudocode can be found in Fig. S16b,c.

### Multi-view spectral embedding

In this section we extend our matrix-free spectral embedding method to perform dimensionality reduction on multi-modal single-cell data. Assume we have data in multiple views, *e*.*g*., chromatin accessibility and gene expressions, represented by a sequence of count matrices 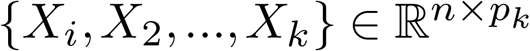. Our objective is to obtain a low-dimensional representation of the data while preserving cell similarity in each view using the spectral embedding method. One approach involves calculating the similarity matrix for each view, normalizing them, and subsequently summing them. The resulting matrix is then used to compute the spectral embedding. This straightforward strategy has proven effective in revealing clusters in prior research^47,48^. However, it necessitates the computation of the similarity matrix for each view, which is computationally demanding. In this section, we present an algorithm that is both time and space-efficient for computing this embedding.

We first normalize each *X*_*i*_ such that the rows of *X*_*i*_ have unit *L*_2_ norm. We then define *X* as the horizontally concatenated view of the sequence of matrices,

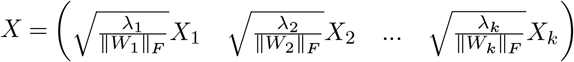

where *λ*_*k*_ is the user defined weights measuring the relative importance of each view; 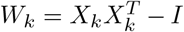 is the similarity matrix of the *k*-th view; ∥*W*_*k*_∥_*F*_ is the Frobenius norm of *W*_*k*_. We can see that,

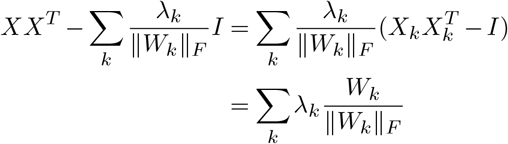

Without loss of generality, we can assume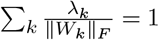. In practice, this can be achieved by normalizing *λ*_*k*_.

The above equation can now be written as,

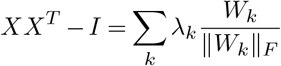

Therefore, the matrix *W* = *XX*^*T*^ −*I* is a linear combination of the normalized similarity matrices of the individual views. To compute the spectral embedding of *W*, it suffices to apply the matrix-free spectral embedding method described above to the concatenated view *X*. This algorithm is implemented in the “snapatac2.tl.multi_spectral” function from the SnapATAC2 package. The pseudocode for this algorithm is shown in Fig. S16d.

### Eigenvector selection in spectral embedding

Not all eigenvectors produced by spectral embedding are informative and relevant for clustering tasks. Selecting appropriate eigenvectors is essential, as using uninformative or irrelevant ones can lead to suboptimal clustering results. We found that the widely-used elbow method for determining the number of eigenvectors is not consistently reliable in practice. To identify relevant eigenvectors, we propose a simple heuristic based on the eigenvalues of the graph Laplacian matrix. In this approach, each eigenvector is weighted by the square root of its corresponding eigenvalue, and these weighted eigenvectors are then employed for further analyses.

### Benchmarking datasets

Synthetic scATAC-seq dataset and the Buenrostro2018 dataset were downloaded from^22^. Additional nine scATAC-seq benchmarking datasets were listed in Table 1. The scHi-C datasets were downloaded from^18^. The scRNA-seq datasets were downloaded from^35^. All benchmarking datasets are converted into a uniform format and are available at https://osf.io/hfs2v/.

### Benchmarking algorithms

In order to ensure the reproducibility of our benchmarking results, we have prepared a separate Docker image for each method. These images can be downloaded from https://hub.docker.com/u/kaizhang. In this section, we provide a brief overview of each method’s usage. To access the code required for reproducing the benchmarking results, visit: https://github.com/kaizhang/single-cell-benchmark.

### ArchR

ArchR is a R package for analyzing scATAC-seq data. To generate the lower-dimensional embedding of the data, we used the “ArchR:::.computeLSI” function with the default parameters. The output dimension was set to 30. After performing the SVD, ArchR scales the singular vectors by the singular values. As a result, component selection is not necessary so we used all 30 dimensions for downstream analysis. Note ArchR includes three variants of the LSI algorithm: “TF-logIDF”, “log(TF-IDF)”, and “logTF-logIDF”. Although we have benchmarked all three variants, we only report the results for the “log(TF-IDF)” variant in the main text as it is the default setting.

### Signac

Signac is a R package for analyzing scATAC-seq data. To generate the lower-dimensional embedding of the data, we used the “Signac:::RunTFIDF.default” and “Signac:::RunSVD.default” functions with the default parameters. The output dimension was set to 30. We used the elbow method to select the number of components for downstream analysis. Note Signac includes four variants of the LSI algorithm: “IDF”, “TF-logIDF”, “log(TF-IDF)”, and “logTF-logIDF”. Although we have benchmarked all four variants, we only report the results for the “log(TF-IDF)” variant in the main text as it is the default setting.

### EpiScanpy

Episcanpy is a Python package for analyzing scATAC-seq data. We first normalized the count matrix using “episcanpy.pp.normalize_per_cell” and “episcanpy.pp.log1p” functions with the default parameters. We then used the “episcanpy.pp.pca” function to generate the lower-dimensional embedding of the data. The output dimension was set to 30.

### SCALE

SCALE is a Python package for performing dimensionality reduction on scATAC-seq data. We used the command “SCALE.py” with following parameters to generate the lower-dimensional embedding: “–min_peaks 0 –min_cells 0 -l 30”. Additionally, as we knew the number of cell types in the benchmarking datasets, we set the “-k” parameter (the number of clusters) to the true number of cell types.

### PeakVI

PeakVI is a Python package for performing dimensionality reduction on scATAC-seq data. We used the “scvi.model.PEAKVI” function to create a model with the default parameters. The dimensionality of the latent variable was set to 30.

### scBasset

scBasset is a Python package for performing dimensionality reduction on scATAC-seq data. We followed the instructions in the scBasset GitHub repository to generate the lower-dimensional embedding of the data. The dimensionality of the latent variable was set to 30.

### pycisTopic

pycisTopic is a Python package for analyzing scATAC-seq data. To generate the lower-dimensional embedding of the data, we first created a model using the “create_cistopic_object” function with the default parameters. We then used “run_cgs_models” to train the model with the following parameters: “n_iter=300, alpha=50, alpha_by_topic=True, eta=0.1, eta_by_topic=False”. We trained six models with different dimensions of the latent variable: 5, 10, 15, 20, 25, and 30. We then used the “evaluate_models” function to select the best model for downstream analysis.

### SnapATAC

SnapATAC is a R package for analyzing scATAC-seq data. For datasets with number of cells less than 20,000, we used the “SnapATAC::runDiffusionMaps” function with the default parameters to generate the lower-dimensional embedding of the data. For datasets with number of cells more than 20,000, running “SnapATAC::runDiffusionMaps on the full dataset requires a large amount of memory. In this case, we applied “SnapATAC::runDiffusionMaps” on a subset of the data and then used the “SnapATAC::runDiffusionMapsExtension” function to generate the lower-dimensional embedding of the full dataset. The output dimension was set to 30. We used the “SnapATAC:::weightDimReduct” to scale eigenvectors by their corresponding eigenvalues. The scaled eigenvectors were then used for downstream analysis.

### Higashi

Higashi is a Python package for analyzing scHi-C data. We followed the instructions in the Higashi GitHub repository to generate the lower-dimensional embedding of the data. The dimensionality of the cell embeddings was set to 30.

### scHiCluster

scHiCluster is a Python package for analyzing scHi-C data. We followed the instructions in the scHi-Cluster GitHub repository to generate the lower-dimensional embedding of the data. The dimensionality of the cell embeddings was set to 30.

### SCANPY

SCANPY is a Python package for analyzing scRNA-seq data. To generate the lower-dimensional embedding of the data, we first applied the “scanpy.pp.normalize_total” and “scanpy.pp.log1p” functions to preprocess the data. We then used the “scanpy.pp.highly_variable_genes” function to select highly variable genes with “n_top_genes=5000”. The data was then scaled using “scanpy.pp.scale” with “max_value=10” and inputed to the “scanpy.tl.pca” function to get lower-dimensional embedding. The output dimension was set to 30.

### Benchmarking metrics

We generated a lower-dimensional embedding for each method and used it to create a *k*-nearest neighbor graph with *k* set at 50. Next, we applied the Leiden algorithm to the *k*-nearest neighbor graph to cluster cells. Since the number of cell types in the benchmarking datasets is known, we adjusted the Leiden algorithm’s resolution parameter from 0.1 to 3.0 in increments of 0.1 to produce the same number of clusters. We utilized the adjusted Rand index (ARI) and adjusted mutual information (AMI) metrics to evaluate the clustering results.

## Data availability

We do not generate any new data in this study. The benchmark datasets used in this study can be accessed at https://osf.io/hfs2v/.

## Code availability

The source code of SnapATAC2 can be accessed at https://github.com/kaizhang/SnapATAC2. The source code for reproducing the benchmarks in this project can be accessed at https://github.com/kaizhang/single-cell-benchmark.

## Acknowledgements

We thank Y. Zhang, S. Zu and Y.E. Li for valuable discussion and feedback in earlier versions of the manuscript. This work was supported by the Ludwig Institute for Cancer Research (to B.R.), and National Human Genome Research Institute (3U54HG006997-04S2 to B.R.).

## Author contributions

Conceptualization: K.Z. and B.R.; Methodology: K.Z.; Software: K.Z.; Investigation: K.Z. and B.R.; Data Curation: K.Z., N.R.Z., and E.J.A.; Writing – Original Draft: K.Z. and B.R.; Writing – Review and Editing: K.Z., N.R.Z., E.J.A. and B.R.; Funding Acquisition: B.R.

## Supplementary Note 1: Supplementary material

**Extended Data Table 1** | **Comparing the Runtime and memory usage of different dimensionality reduction methods**.

**Extended Data Table 2** | **Comparing the runtime and memory usage of the preprocessing components in ArchR and SnapATAC2**.

**Extended Data Table 3** | **Benchmarking the cell embedding methods on scATAC-seq datasets. Extended Data Table 4** | **Benchmarking the cell embedding methods on scHi-C datasets**.

**Extended Data Table 5** | **Benchmarking the cell embedding methods on scRNA-seq datasets**.

**Extended Data Table 6** | **Comparing the cell embeddings generated by the Nystrom method with those produced using the subsampled LSI method**.

**Figure S1.**
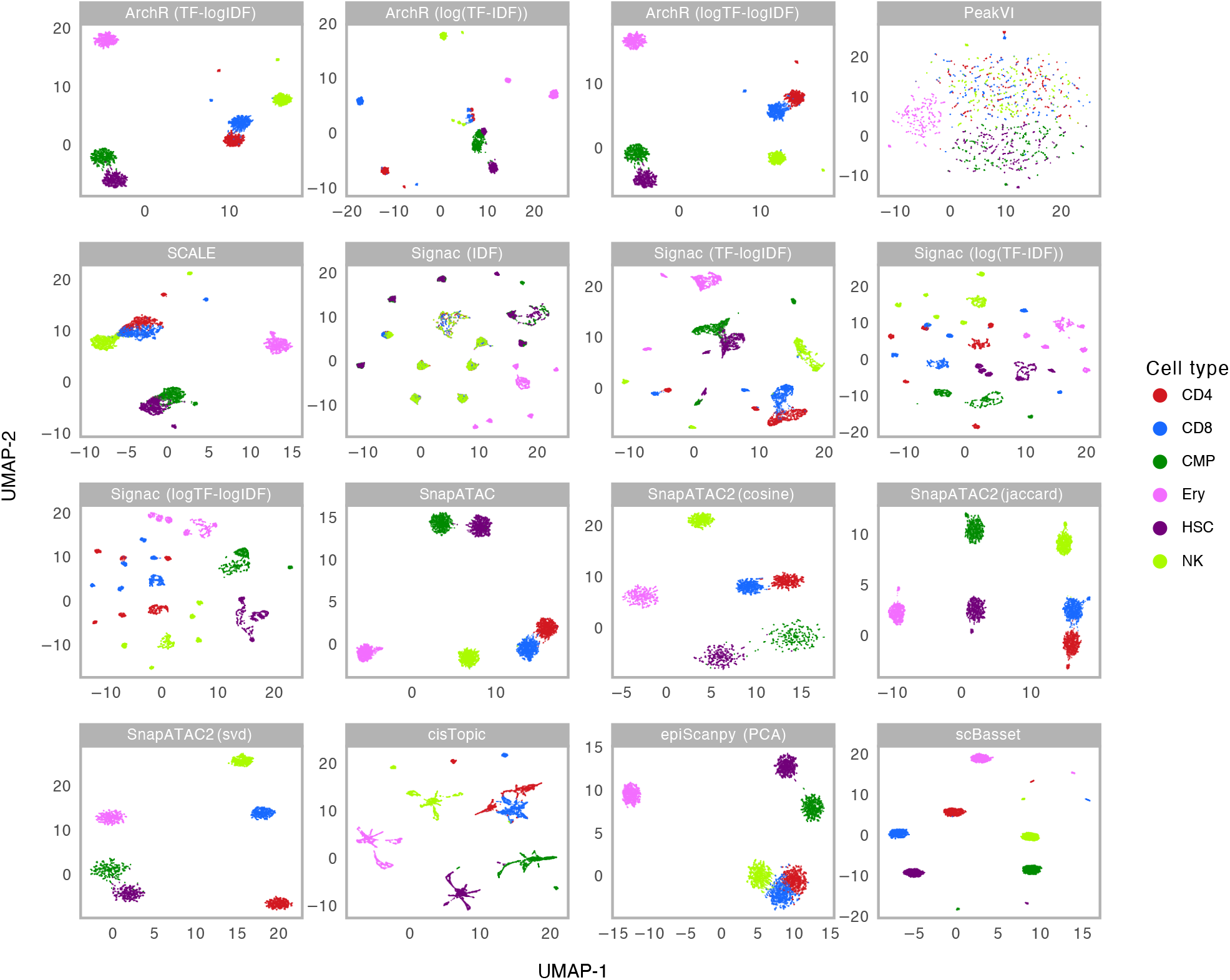
Cell embeddings for the Chen *et al*. dataset. UMAP visualization of different cell embedding methods on the dataset from Chen *et al*. Cells are color-coded by cell type labels. ArchR (TF-logIDF): the embeddings produced by ArchR using the “TF-logIDF” method. ArchR (log(TF-IDF)): the embeddings produced by ArchR using the “log(TF-IDF)” method. This is the default method used by ArchR and was reported in the main text. ArchR (logTF-logIDF): the embeddings produced by ArchR using the “logTF-logIDF” method. PeakVI: the embeddings produced by PeakVI using the variational autoencoder model. SCALE: the embeddings produced by SCALE. Signac (IDF): the embeddings produced by Signac using the “IDF” method. Signac (TF-logIDF): the embeddings produced by Signac using the “TF-logIDF” method. Signac (log(TF-IDF)): the embeddings produced by Signac using the “log(TF-IDF)” method. This is the default method used by Signac and was reported in the main text. Signac (logTF-logIDF): the embeddings produced by Signac using the “logTF-logIDF” method. SnapATAC: the embeddings produced by SnapATAC using the spectral embedding with Jaccard index as the similarity measure. SnapATAC2 (cosine): the embeddings produced by SnapATAC2 using the spectral embedding with cosine similarity as the similarity measure. SnapATAC2 (Jaccard): the embeddings produced by SnapATAC2 using the spectral embedding with Jaccard index as the similarity measure. SnapATAC2 (svd): Same as “SnapATAC2 (cosine)” except that the diagonal of the similarity matrix is not set to zero. cisTopic: the embeddings produced by cisTopic using the latent Dirichlet allocation model. epiScanpy: the embeddings produced by EpiScanpy using the principal component analysis. scBasset: the embeddings produced by scBasset using the convolutional neural network model.

**Figure S2.**
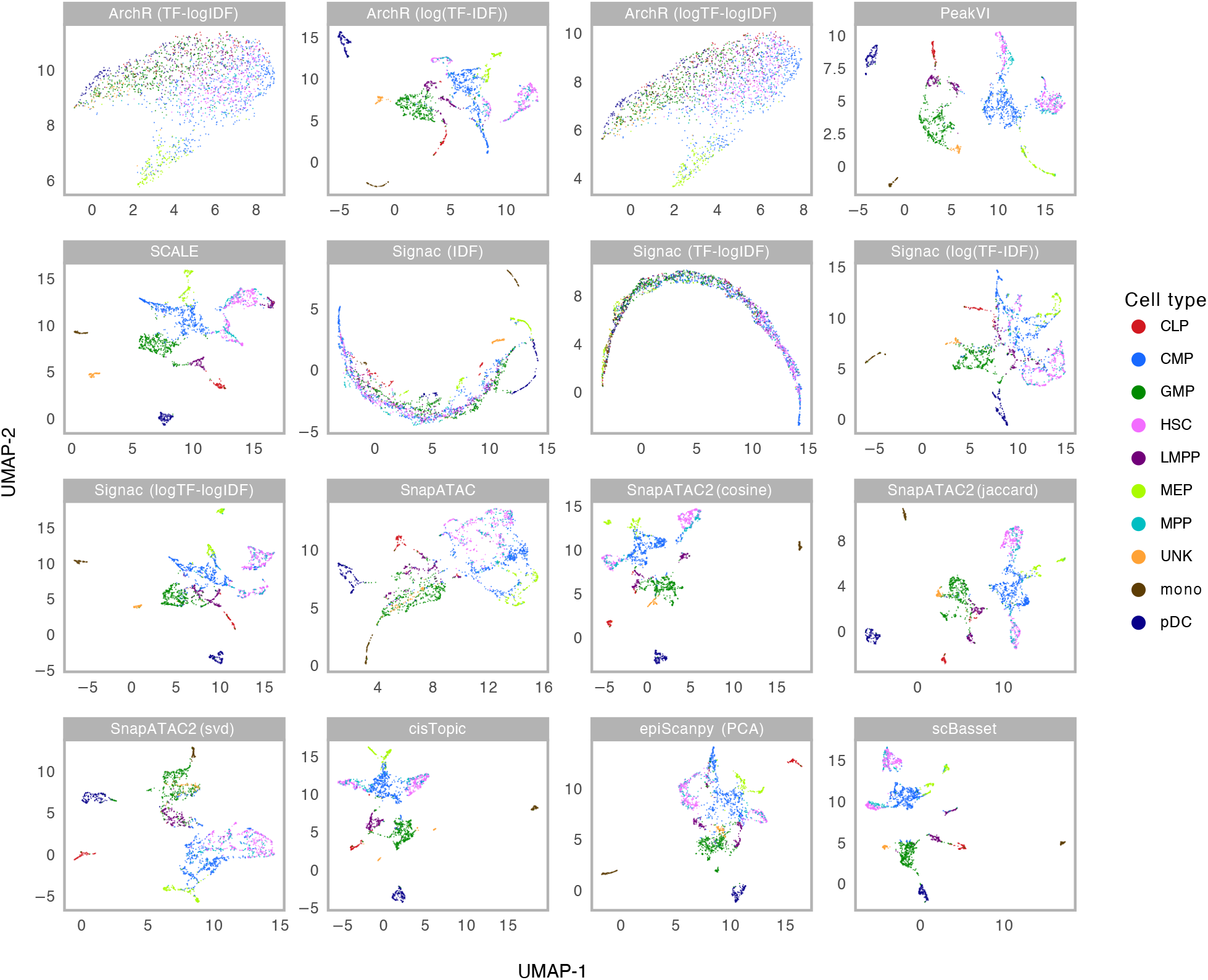
Cell embeddings for the Buenrostro *et al*. dataset. UMAP visualization of different cell embedding methods on the dataset from Buenrostro *et al*. Cells are color-coded by cell type labels. See the caption of **Fig. S1** for the description of the methods.

**Figure S3.**
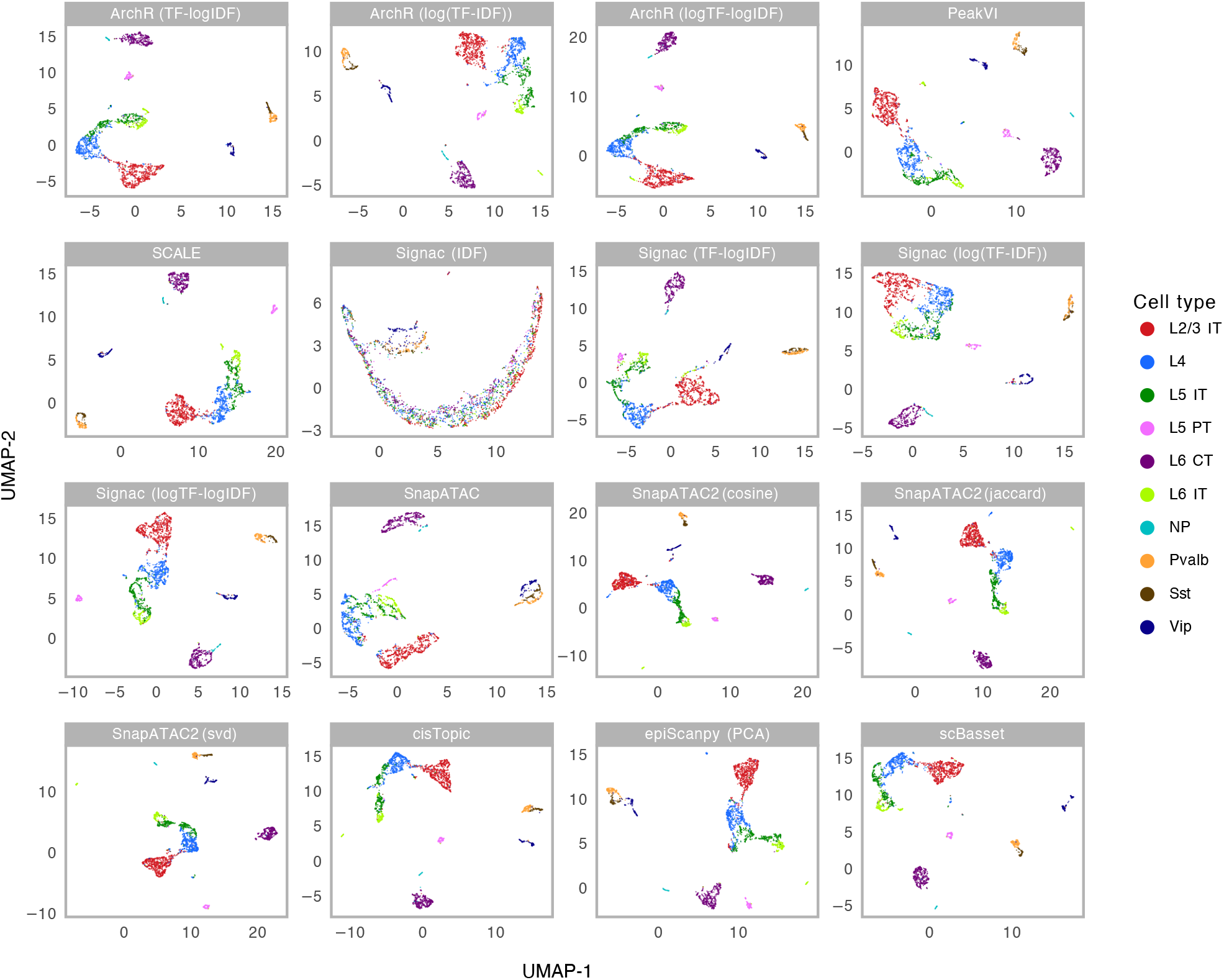
Cell embeddings for the 10x Brain 5k dataset. UMAP visualization of different cell embedding methods on 10x Brain 5k dataset. Cells are color-coded by cell type labels. See the caption of **Fig. S1** for the description of the methods.

**Figure S4.**
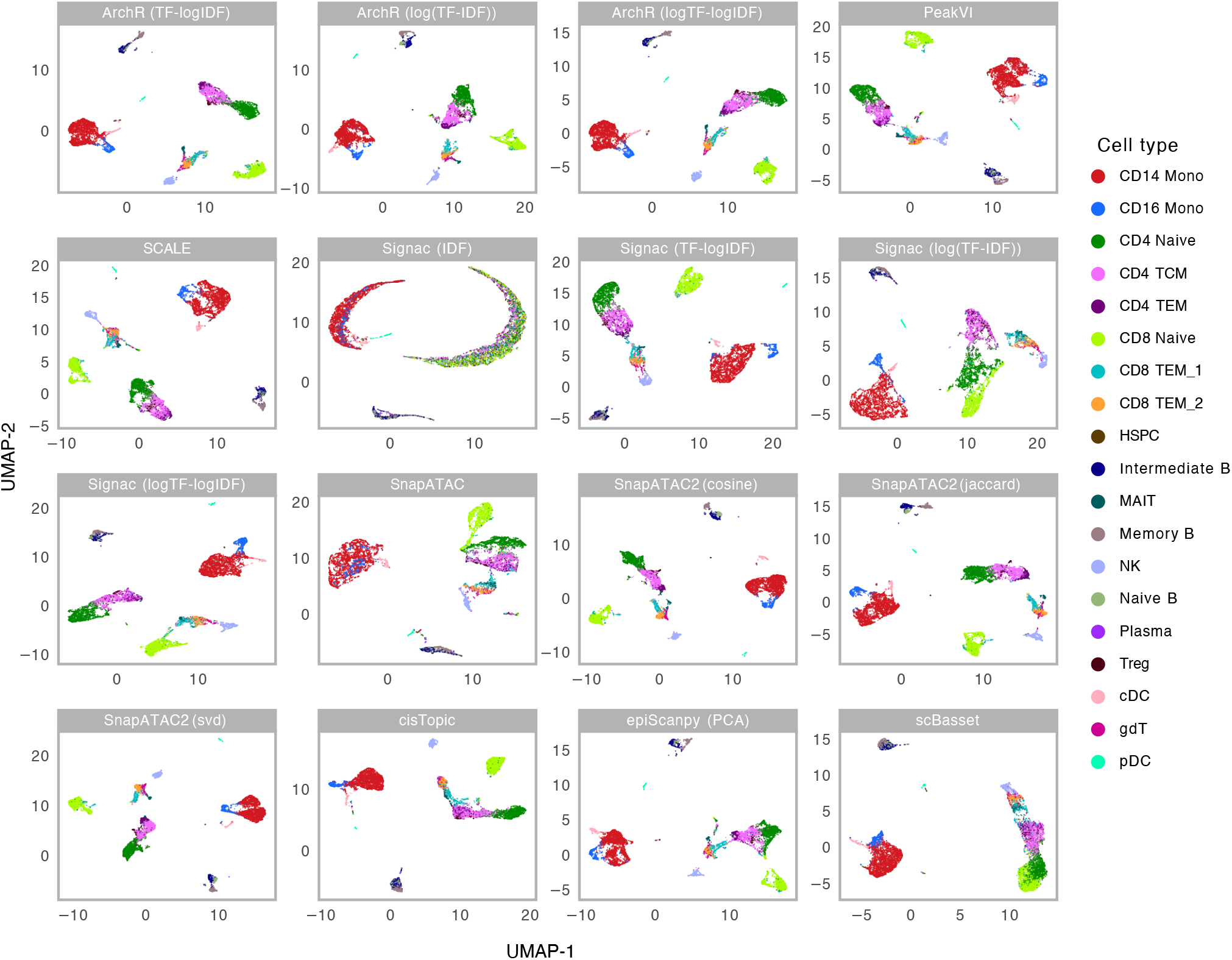
Cell embeddings for the 10x PBMC 10k dataset. UMAP visualization of different cell embedding methods on 10x PBMC 10k dataset. Cells are color-coded by cell type labels. See the caption of **Fig. S1** for the description of the methods.

**Figure S5.**
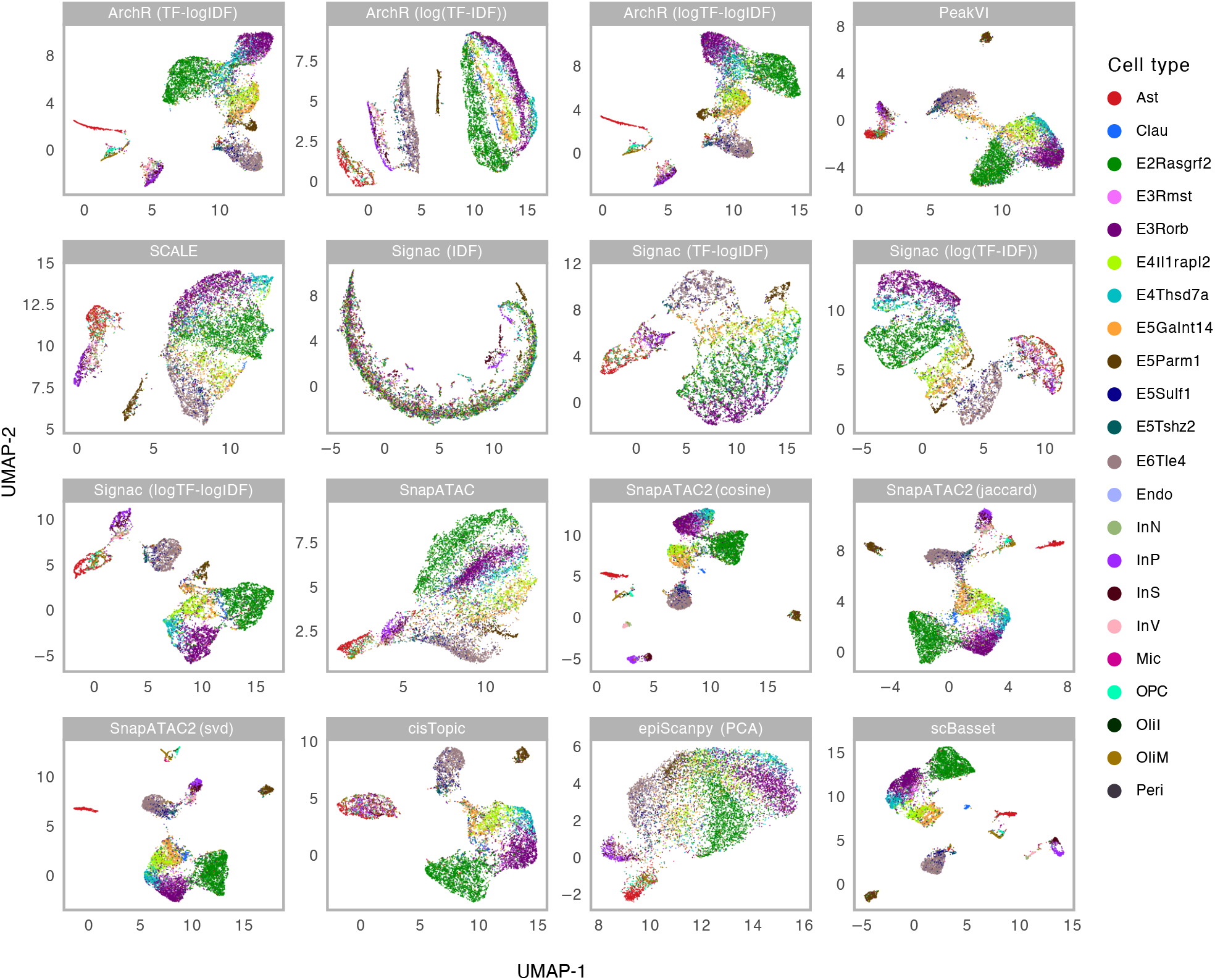
Cell embeddings for the Chen *et al*. dataset. UMAP visualization of different cell embedding methods on the dataset from Chen *et al*. Cells are color-coded by cell type labels. See the caption of **Fig. S1** for the description of the methods.

**Figure S6.**
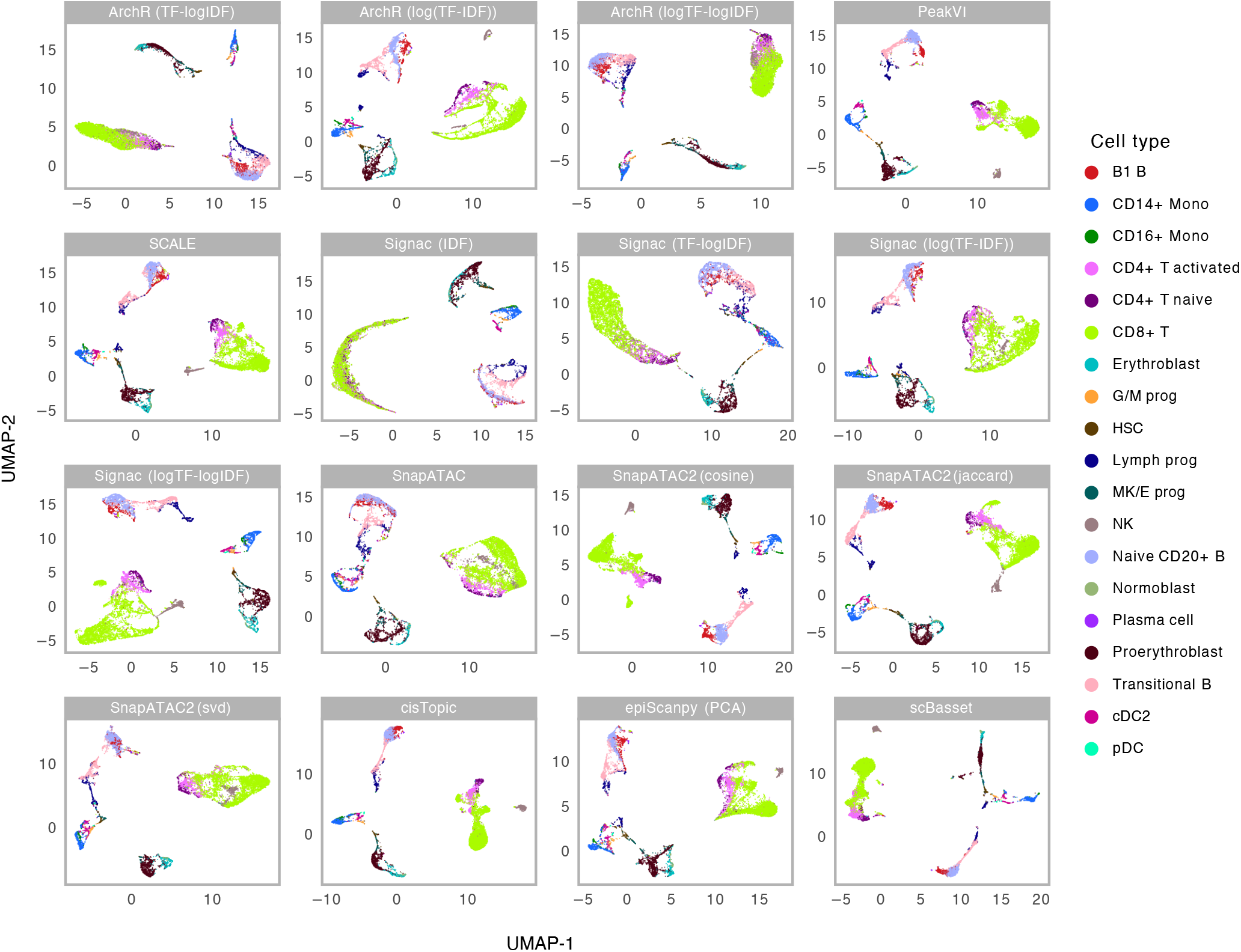
Cell embeddings for the GSE194122 dataset. UMAP visualization of different cell embedding methods on the dataset from GSE194122. Cells are color-coded by cell type labels. See the caption of **Fig. S1** for the description of the methods.

**Figure S7.**
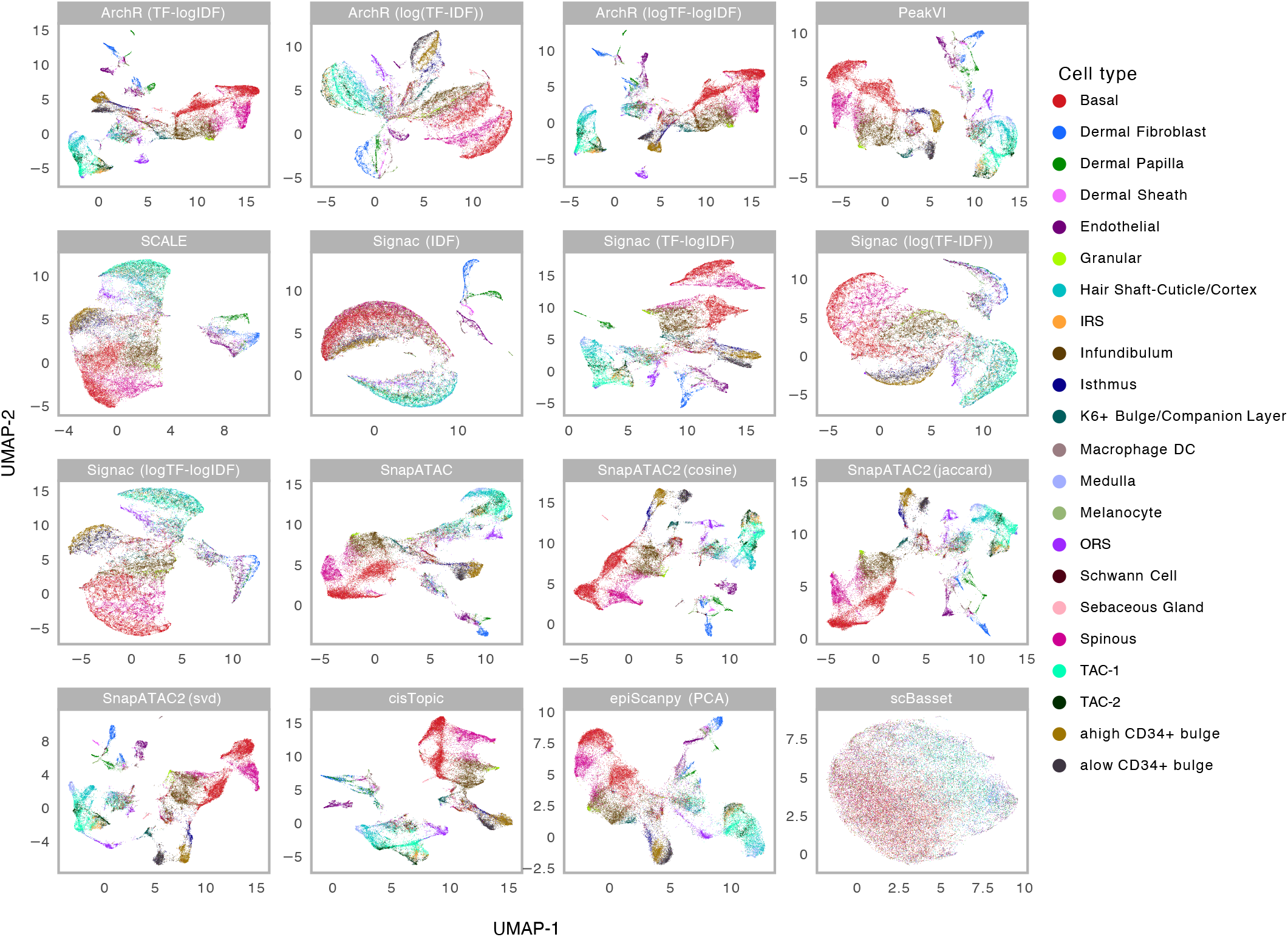
Cell embeddings for the Ma *et al*. dataset. UMAP visualization of different cell embedding methods on the dataset from Ma *et al*. Cells are color-coded by cell type labels. See the caption of **Fig. S1** for the description of the methods.

**Figure S8.**
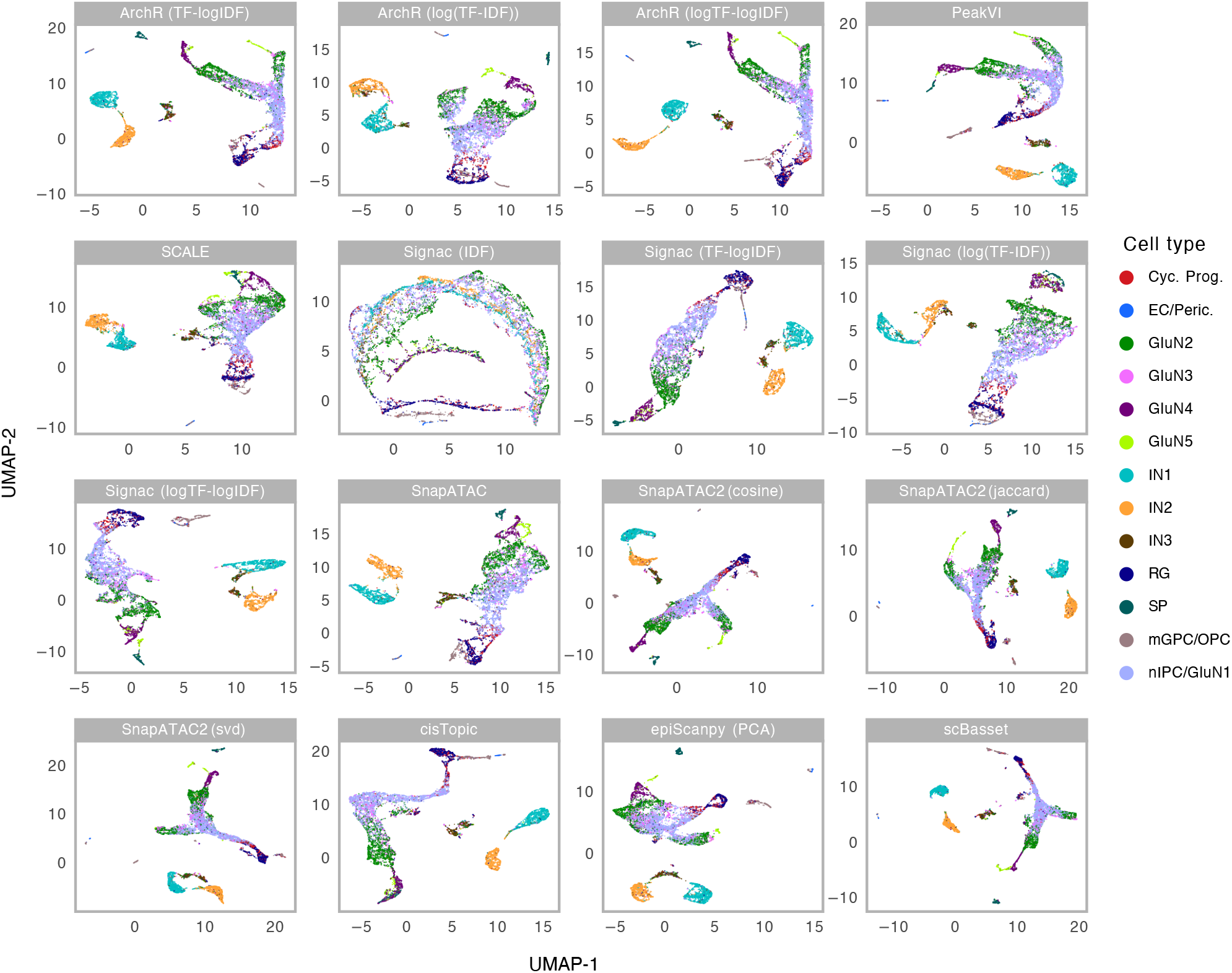
Cell embeddings for the Trevino *et al*. dataset. UMAP visualization of different cell embedding methods on the dataset from Trevino *et al*. Cells are color-coded by cell type labels. See the caption of **Fig. S1** for the description of the methods.

**Figure S9.**
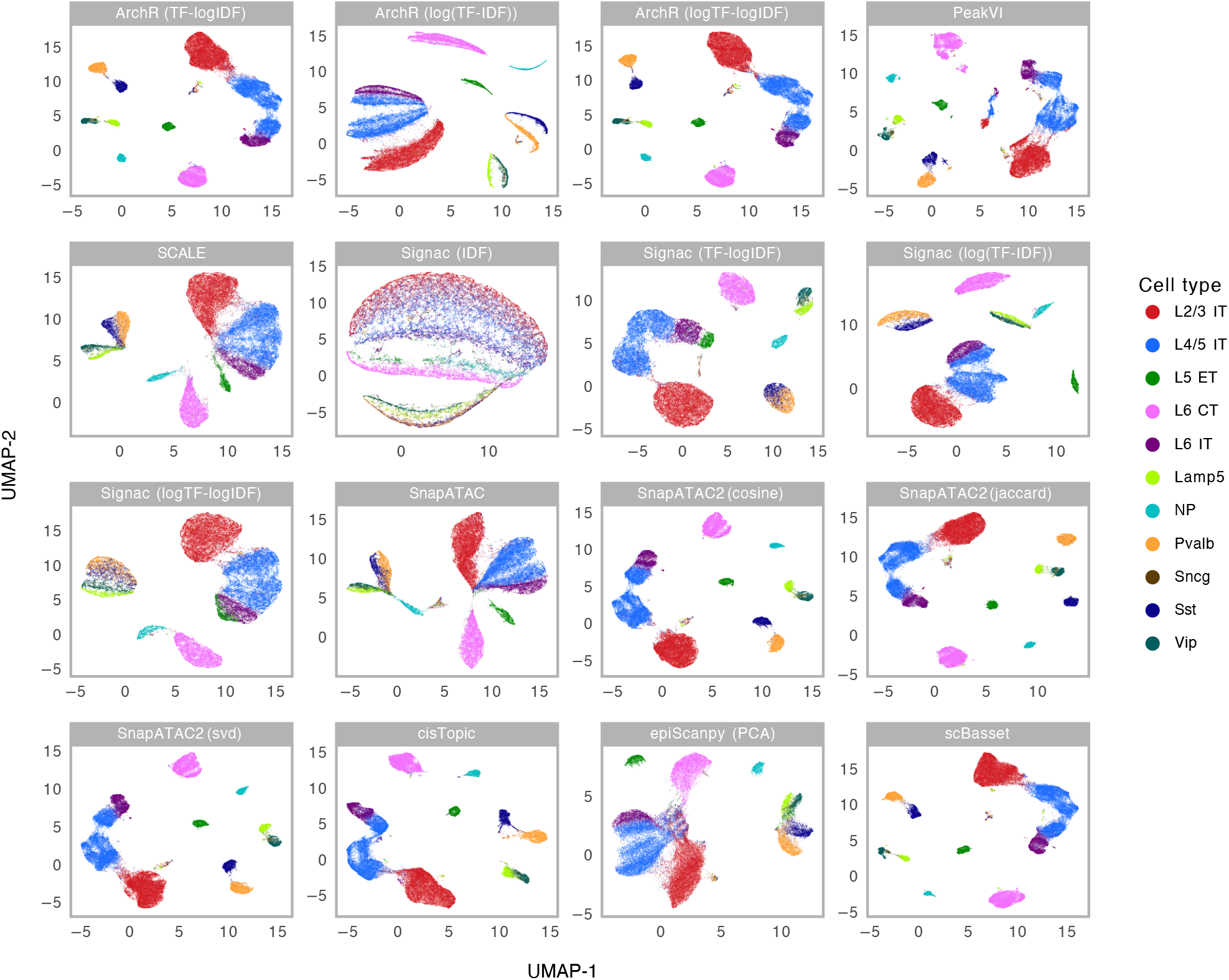
Cell embeddings for the Yao *et al*. dataset. UMAP visualization of different cell embedding methods on the dataset from Yao *et al*. Cells are color-coded by cell type labels. See the caption of **Fig. S1** for the description of the methods.

**Figure S10.**
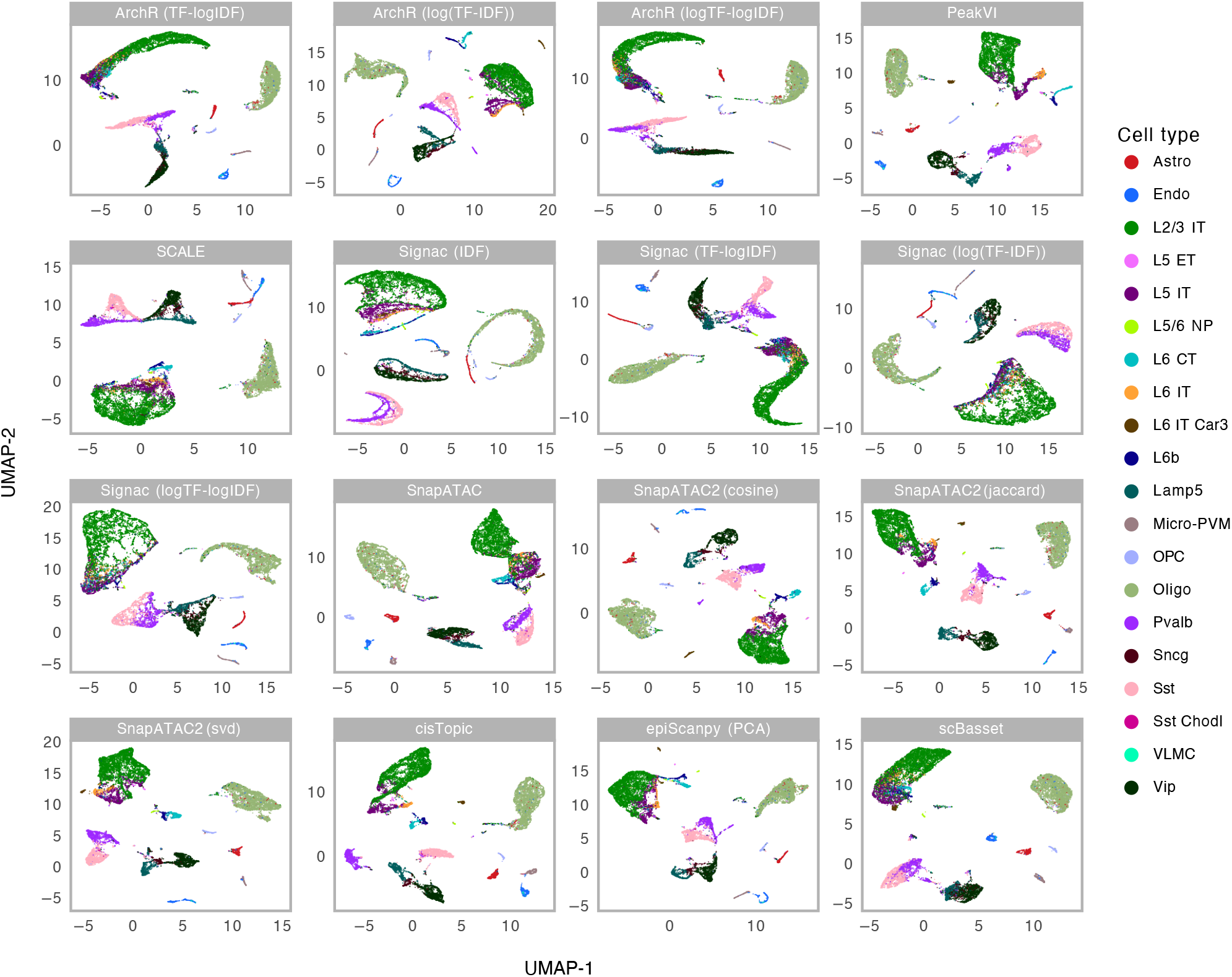
Cell embeddings for the Zemke *et al*. human dataset. UMAP visualization of different cell embedding methods on the human dataset from Zemke *et al*. Cells are color-coded by cell type labels. See the caption of **Fig. S1** for the description of the methods.

**Figure S11.**
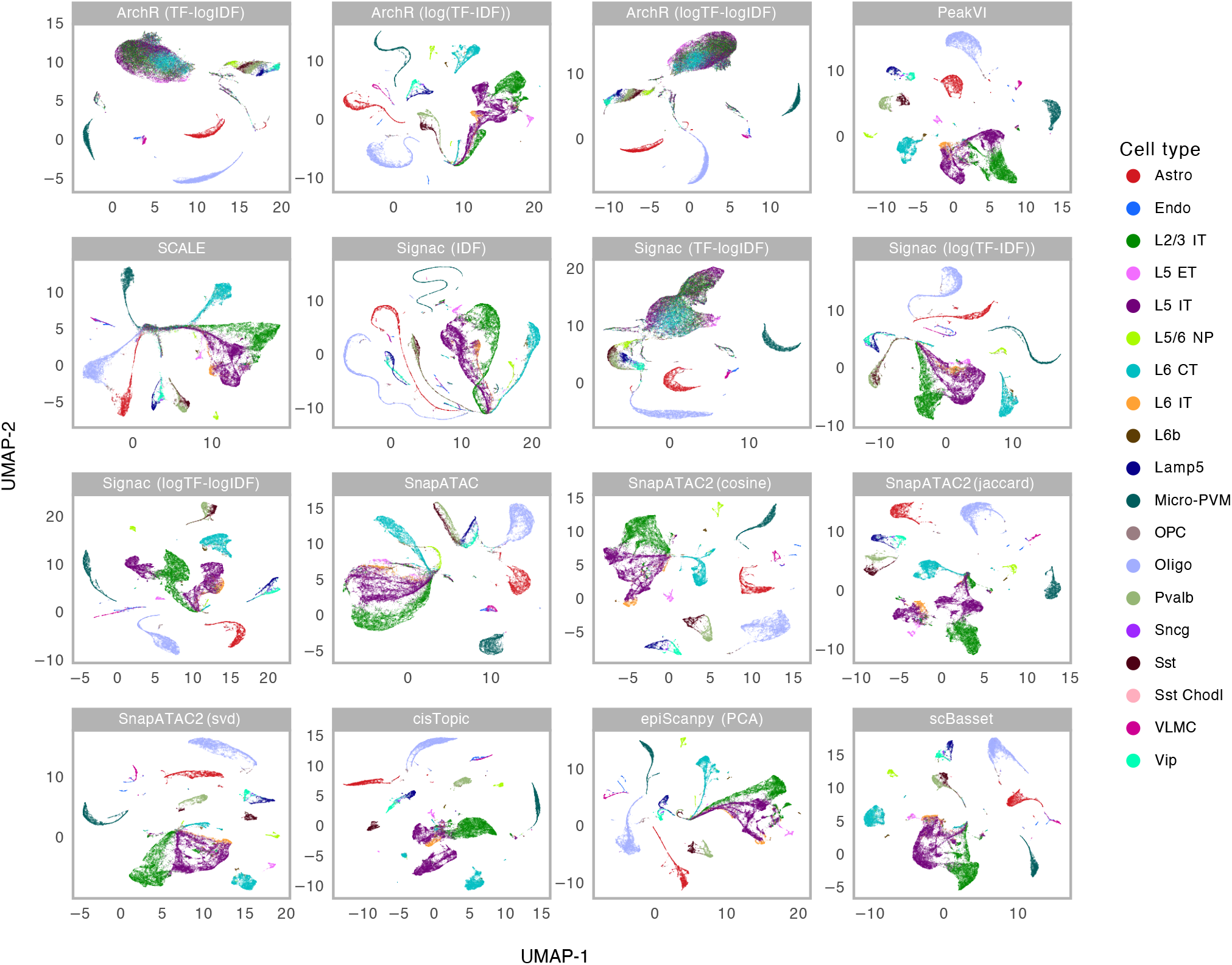
Cell embeddings for the Zemke *et al*. mouse dataset. UMAP visualization of different cell embedding methods on the mouse dataset from Zemke *et al*. Cells are color-coded by cell type labels. See the caption of **Fig. S1** for the description of the methods.

**Figure S12.**
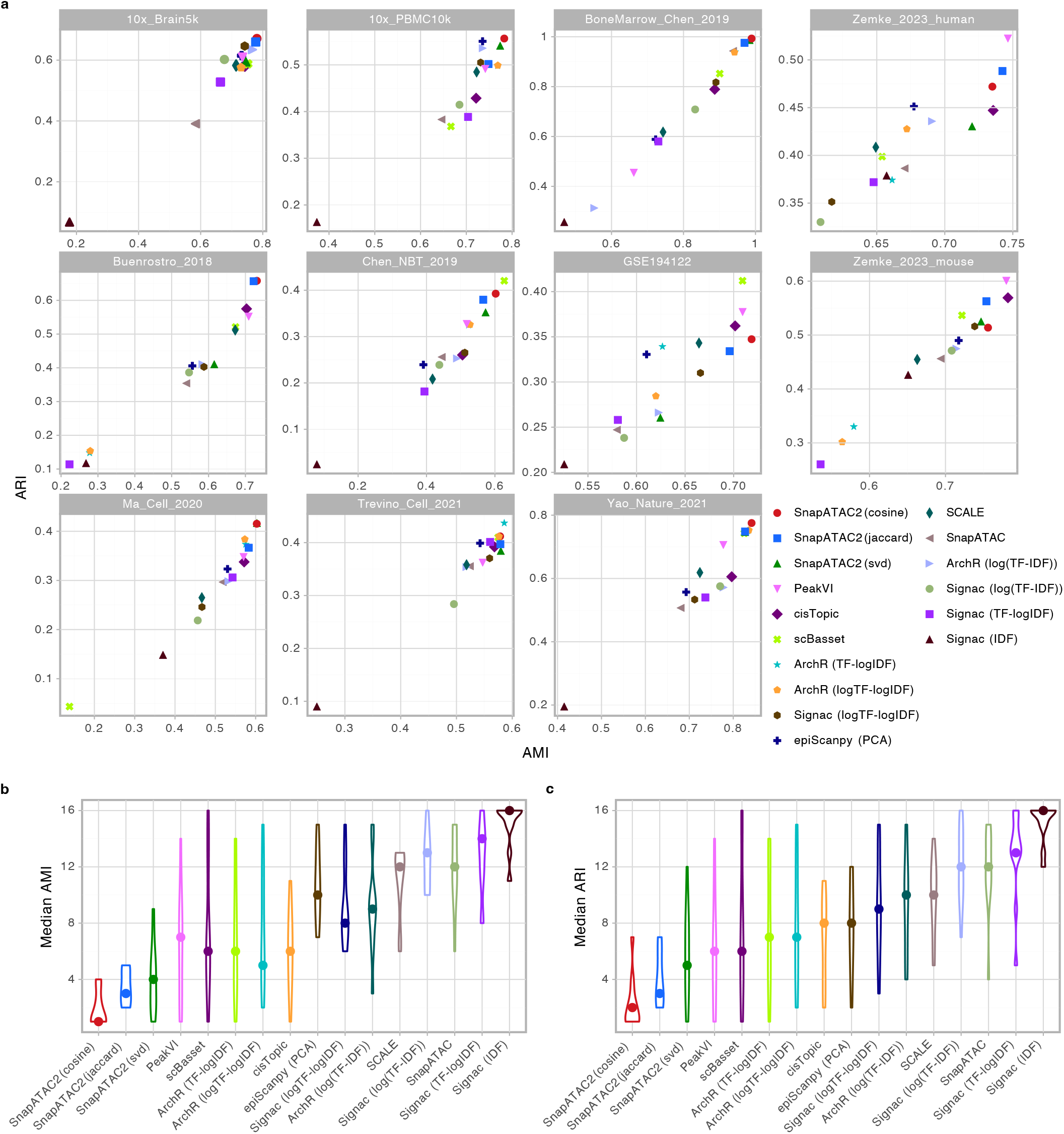
Summary of benchmarking results of various dimensionality reduction methods on scATAC-seq datasets. **a**, Scatter plots showing the relationship between the adjusted mutual information (AMI) and the adjusted Rand index (ARI) for the cell embeddings produced by different methods. The AMI and ARI are computed between the cell type labels and the clusters identified by each method. See the caption of **Fig. S1** for the description of the methods. **b**,**c**, Violin plots showing the distribution of AMI (**b**) and ARI (**c**) scores for each method across all eleven scATAC-seq benchmark datasets. The median rank of each method is indicated by the dot in the violin plot.

**Figure S13.**
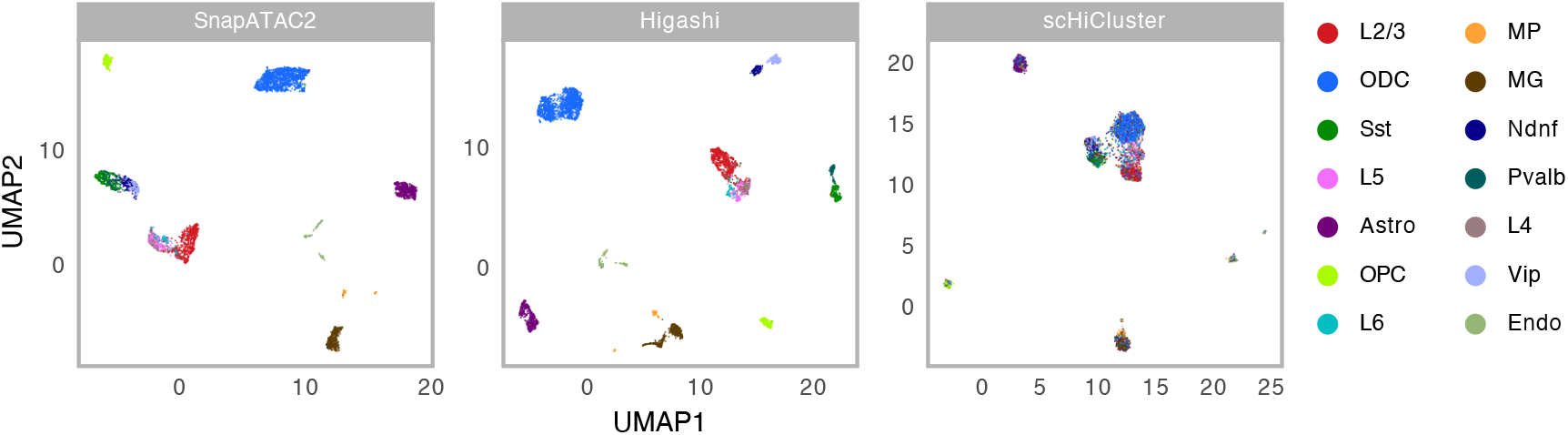
UMAP visualization of the embeddings produced by SnapATAC2, Higashi, and scHiCluster on single-cell Hi-C dataset from Lee et al.

**Figure S14.**
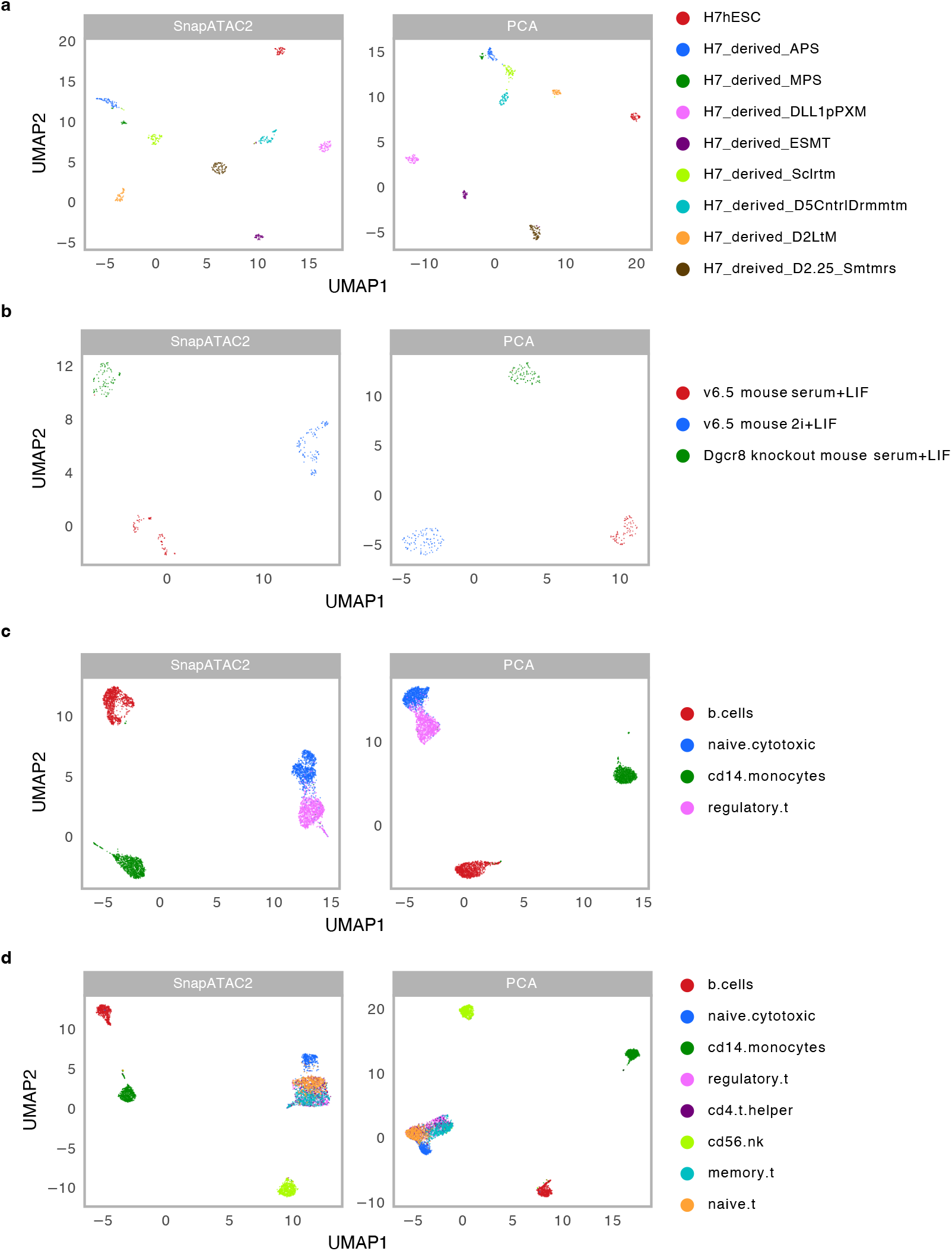
UMAP visualization of the embeddings produced by SnapATAC2 and PCA (scanpy) on various single cell RNA-seq datasets. The datasets include Koh (**a**), Kumar (**b**), Zhengmix4eq (**c**), Zhengmix8eq (**d**).

**Figure S15.**
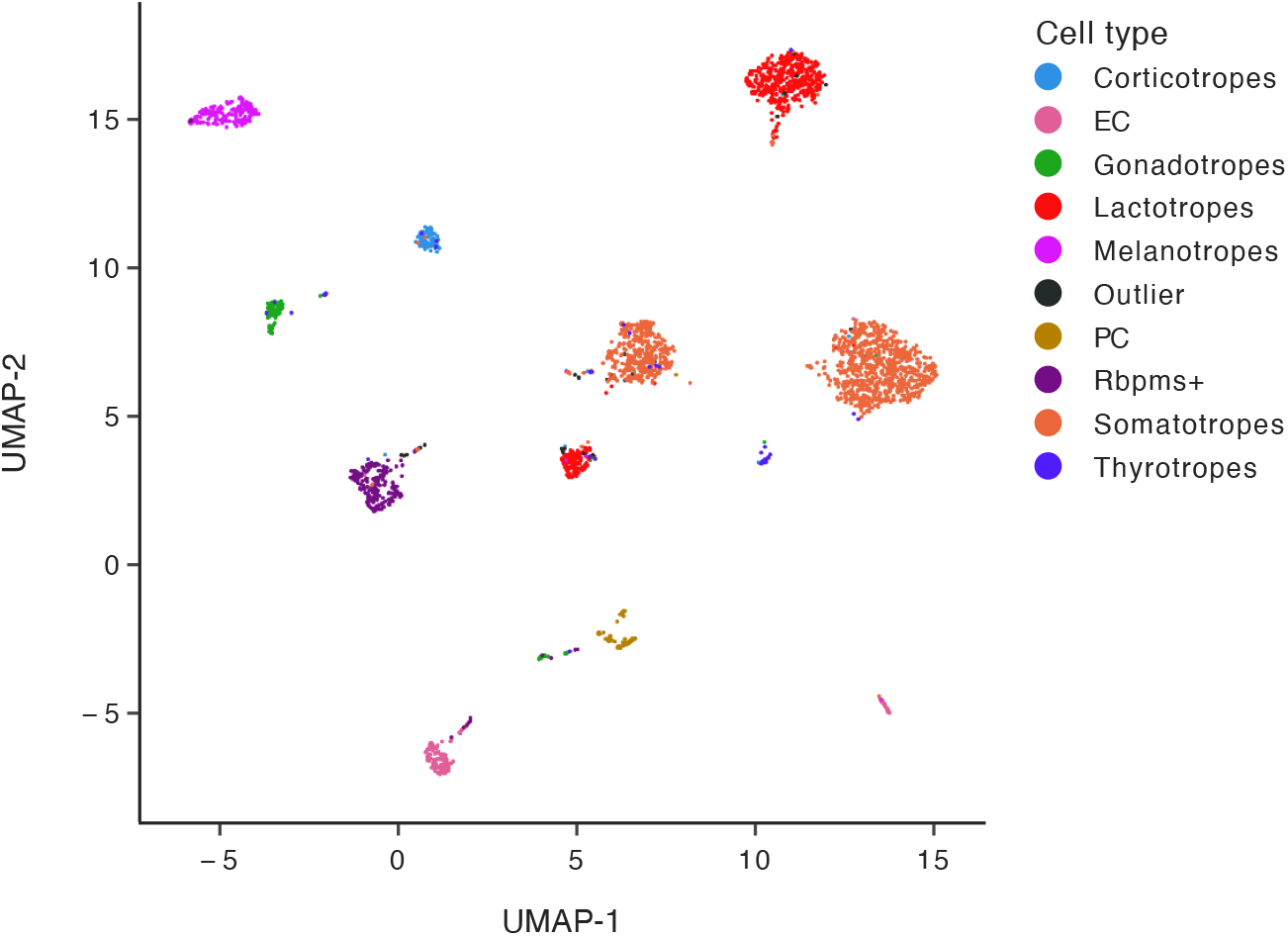
SnapATAC2 unveils fine-grained cellular heterogeneity in single-cell DNA methylation data from Ruf-Zamojski *et al*.

**Figure S16.**
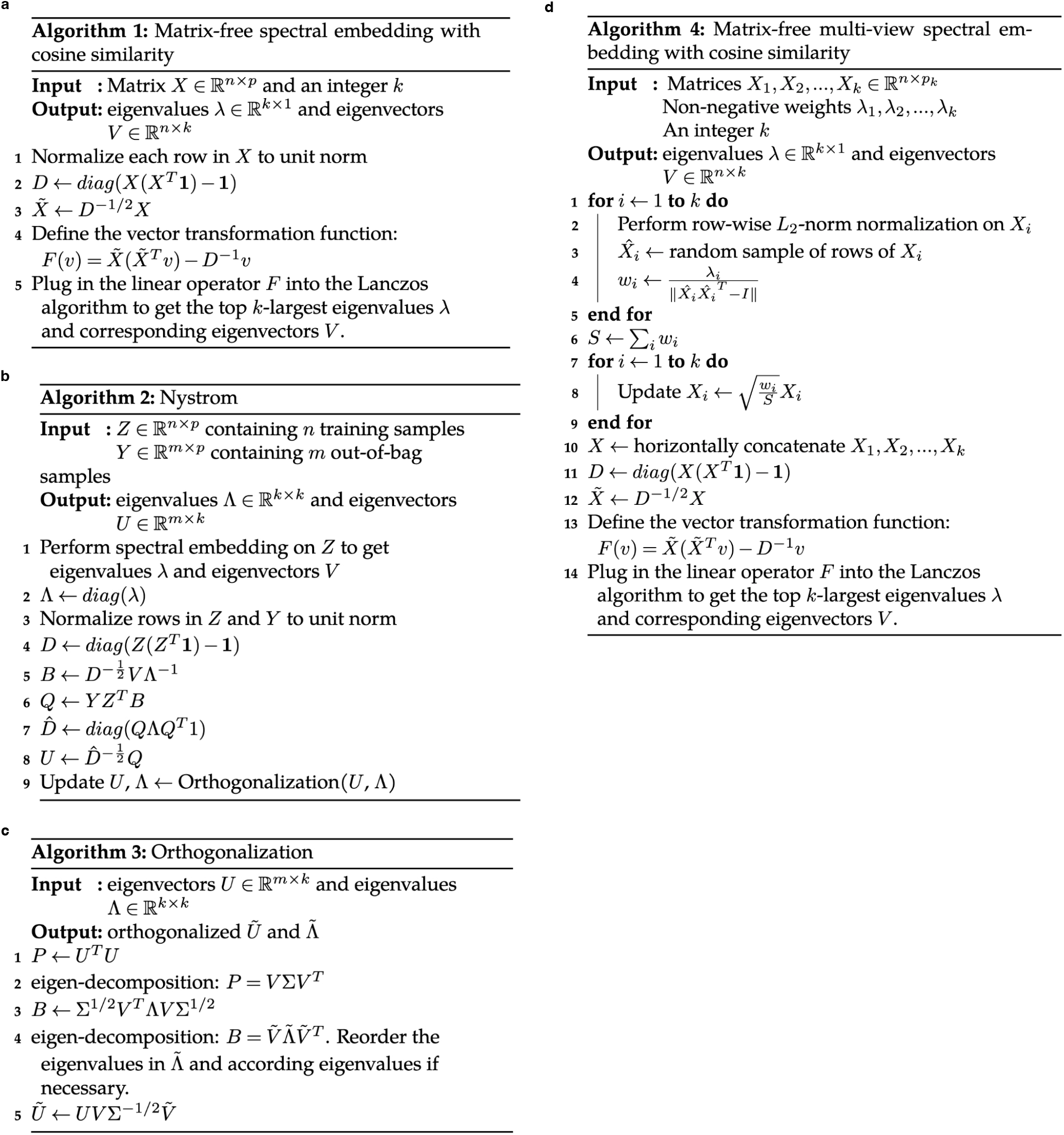
The pseudocodes of various algorithms used in this study. **a**, The pseudocode of the matrix-free spectral embedding algorithm. **b**, The pseudocode of the Nyström algorithm for performing the out-of-sample embedding. **c**, The pseudocode for performing orthogonalization on the eigenvectors produced by the Nyström algorithm. **d**, The pseudocode of the matrix-free multi-view spectral embedding algorithm.

